# Inhibition of the RNA Regulator HuR mitigates spinal cord injury by potently suppressing post-injury neuroinflammation

**DOI:** 10.1101/2024.10.25.620341

**Authors:** Mohammed Amir Husain, Reed Smith, Robert E. Sorge, Abdulraheem Kaimari, Ying Si, Ali Z. Hassan, Abhishek Guha, Katherine A. Smith, Christopher P. Cardozo, Jennifer J. DeBerry, Shaida A. Andrabi, L. Burt Nabors, Natalia Filippova, Caroline K. Webb, Peter H. King

## Abstract

**Background:** Neuroinflammation plays a significant role in promoting secondary tissue injury after spinal cord trauma. Within minutes after spinal cord injury (SCI), microglia and astrocytes become activated and produce inflammatory mediators such as TNF-α, IL-6, iNOS and COX-2 which induce tissue injury through cytotoxicity, vascular hyperpermeability, and secondary ischemia. The inflammatory cascade is amplified by chemokines such as CCL2 and CXCL1 that promote recruitment of peripheral inflammatory cells into the injured spinal cord. HuR is a key post-transcriptional RNA regulator that controls glial expression of many pro-inflammatory factors by binding to adenylate- and uridylate-rich elements in 3’ untranslated regions of the mRNA. SRI-42127 is a small molecule inhibitor that blocks HuR nucleocytoplasmic translocation, a process critical for its regulatory function. The goal of this study was to assess the potential of SRI-42127 for suppressing neuroinflammation after SCI and improving functional outcome.

**Methods:** Adult female mice underwent a contusion injury at the T10 level. SRI-42127 or vehicle was administered intraperitoneally starting 1 h after injury and up to 5 days. Locomotor function was assessed by open field testing, balance beam and rotarod. Immunohistochemistry was used to assess lesion size, neuronal loss, myelin sparing, microglial activation and HuR localization. Molecular analyses of spinal cord and peripheral tissues for expression of inflammatory mediators included qPCR, immunohistochemistry, ELISA, or western blot. Post-SCI pain was assessed by the mouse grimace scale.

**Results:** SRI-42127 significantly attenuated loss of locomotor function and post-SCI pain. Histologic correlates to these beneficial effects included reduced lesion size, neuronal loss, and an increase in myelin sparing. There was reduced microglial activation at the epicenter with concomitant attenuation of HuR nucleocytoplasmic translocation. Molecular analysis revealed a striking reduction of pro-inflammatory mediators at the epicenter including IL-6, MMP-12, IL-1β, TNF-α, iNOS, COX-2, and chemokines CCL2, CXCL1, and CXCL2. Suppression of inflammatory responses extended peripherally including serum, liver, and spleen.

**Conclusion:** Targeting HuR after SCI is a viable therapeutic approach for suppressing neuroinflammatory responses after tissue injury and improving functional outcome.

## Background

Traumatic spinal cord injury (SCI) is a debilitating neurological injury affecting more the 250,000 individuals per year globally. The injury often results in in permanent disability and shortened lifespan. Additionally, there are life-changing socioeconomic consequences for both the individual and their caregivers, particularly younger people [1, 2]. Treatment in the acute phase of injury is mainly supportive with limited treatment options for neuroprotection to improve recovery and reduce long term disability.

The physical trauma of SCI triggers a robust inflammatory response initiated within minutes by resident microglia and astrocytes in the vicinity of tissue injury. This activation leads to production of pro-inflammatory mediators including IL-1β, IL-6, TNF-α, MMPs, COX-2 and nitric oxide (NO) via iNOS that exert direct toxic effects on neurons, oligodendroglia, and endothelial cells, often leading to apoptosis and peaking within the initial 24 h post injury [3, 4]. These mediators increase vascular permeability by disrupting the blood-spinal cord barrier (BSCB), leading to edema, secondary ischemia, and hemorrhage [5, 6]. Disruption of the BSCB, in combination with glial secretion of chemokines such as CCL2 and CXCL1, further intensifies the inflammatory cascade by facilitating recruitment and infiltration of peripheral neutrophils and monocytes [3].

An integral regulatory component to this inflammatory cascade, beginning with activated microglia and astrocytes, is at the posttranscriptional level, where AU-rich elements (ARE) in the 3’ untranslated region (UTR) of pro-inflammatory mRNAs govern their expression through modulation of mRNA stability and translational efficiency [4]. HuR, a major cellular RNA-binding protein (RBP) in glial cells, binds to these AREs and positively regulates the stability and translational efficiency leading to increased expression [4, 7–10]. Primarily located in the nucleus, HuR translocates to the cytoplasm upon activation where it transports and stabilizes bound mRNAs and facilitates their localization to polysomes for translation. In microglia and/or astrocytes, HuR translocation is triggered in different inflammatory conditions, including SCI [9, 11], amyotrophic lateral sclerosis [8], lipopolysaccharide (LPS) stimulation [10, 12], hypoxia [10], and glioblastoma [7]. Knockdown or deletion of HuR in microglia or astrocytes significantly diminishes the expression of many key inflammatory mediators, including pro-inflammatory cytokines, chemokines, and iNOS, associated with secondary tissue injury in SCI [7–9]. Our group has recently developed a small molecule, SRI-42127, that blocks HuR dimerization, a process critical for cytoplasmic shuttling and RNA binding [13–15]. SRI-42127 inhibits translocation of HuR in microglial cells and astrocytes after activation with LPS and suppresses the production of pro-inflammatory mediators [10]. In our prior work, SRI-42127 potently attenuates development of neuropathic pain in a peripheral nerve injury model, a process that is driven by neuroinflammatory responses in the spinal cord [16, 17]. With this background, we hypothesized here that inhibition of HuR with SRI-42127 would suppress neuroinflammatory responses after acute SCI, reduce secondary tissue injury, and mitigate loss of motor function.

## Methods

### Animals and contusion model of spinal cord injury

All animal procedures were approved by the UAB Institutional Animal Care and Use Committee and were carried out in accordance with relevant guidelines and regulations of the National Research Council Guide for the Care and Use of Laboratory Animals. Female C57/Bl6 mice between 10–14 weeks of age were anesthetized under isoflurane and a laminectomy was performed to remove the dorsal aspect of the T10 vertebrae. The animal was transferred to a spinal stereotaxic frame and clamps were attached to the T9 and T11 vertebral spines to secure the vertebral column. Using a 0.8-mm diameter tip, a single impact contusion of 50 kdyn was delivered using the Infinite Horizon spinal cord injury device (Precision Systems & Instrumentation, Lexington, KY). Sham control mice received a laminectomy only. Post-operatively, animals received manual bladder evacuation twice daily to prevent urinary tract infections. One dose of subcutaneous buprenorphine was administered prior to injury and every 12 h for 3 d post injury. For molecular studies, SRI-42127 (10 mg/kg) and vehicle were prepared as previously described [16] and administered intraperitoneally every 2 h beginning 1 h after injury for 4 doses. The cohort for behavioral studies received 4 doses daily, starting 1 h after injury, for 5 days.

### RNA isolation and qPCR

The thoracic spinal column was harvested, and 2 mm sections were cut from the epicenter of injury and adjacent rostral, caudal spinal column. Spinal cord was removed from these sections and RNA was isolated using TRIzol reagent as per manufacturer’s protocol (Thermo Fisher Scientific). RNA was precipitated using isopropanol (Sigma) and glycogen (Thermo Fisher Scientific) and concentrations were measured by Nanodrop (Thermo Scientific, Waltham, MA). Complementary DNA was reverse transcribed from 2 μg of total RNA using a reverse transcription kit (Thermo Fisher Scientific). Quantitative Real Time PCR (qRT-PCR) was conducted using the ViiA7-Real-Time PCR system and primers and probes from Applied Biosystems as previously described [10].

### Western blot and ELISA

For western blotting, whole cells lysates were prepared using T-PER (Thermofisher) and quantification was done with a BCA protein assay kit (Thermofisher). Fifty μg of protein per sample were electrophoresed in 4%–20% mini-PROTEAN gel (Biorad) and transferred to a nitrocellulose membrane. Blots were immunostained with a COX-2 antibody (Santa Cruz Biotechnology, USA Inc) was used at a dilution of 1:1000. Each sample was analyzed in triplicate and quantified using Image lab software (Bio-Rad Laboratories, USA Inc.). For ELISA of peripheral serum samples, cytokines (IL-1β, IL-6, IL-10, TNF-α, CCL2, CXCL1) were measured using U-plex biomarker Group 1 (mouse) Assays (K15069M, MSD). Analyses were done using a QuickPlex SQ 120 instrument (MSD) and DISCOVERY WORKBENCH® 4.0 software (MSD).

### Immunohistochemistry

Mice were euthanized 8 h or 3 weeks and perfused with an ice-cold phosphate buffer saline (PBS) solution for 2 min followed by 4% paraformaldehyde (PFA) solution for 5 min [9]. The entire spinal column was then extracted and post-fixed in PFA for 24 h before being decalcified with 8% hydrochloric acid and 10% ethanol in PBS. The spinal column was incubated in a PBS solution with 10% sucrose for cryoprotection then transferred to a 30% sucrose in PBS solution for 48 h. Two mm samples of the spinal column from epicenter and adjacent rostral (+ 2 mm) and caudal tissues (−2 mm) were preserved and cryo-molded in OCT. A cryostat was used to cut samples into 30 μm serial sections cut transversely.

Sections were washed with PBS and blocked in 10% goat serum for 1 h. Samples were incubated overnight with primary antibodies (GFAP, 1:500, Z0334, Dako, Carpinteria, CA), (Iba1, 1:500, FUJIFILM Wako Chemicals U.S.A. Corporation), (NeuN, 1:400, ABN78, Sigma-Aldrich, USA Ltd.), (HuR, 1:500, sc-5261, Santa Cruz Biotechnology, USA Inc). The following day, slide sections were washed and incubated with Alexa fluorophore-conjugated secondary antibodies (Goat anti mouse, A28175, Thermo Fisher Scientific and Goat anti rabbit Cy3, 111-165-144, Jackson Immuno Research) for 2 h, mounted with antifade DAPI + mountant (ThermoFisher Scientific P36966). For myelin staining, sections were incubated with FluoroMyelin Green for 20 minutes (1:300, Invitrogen).

Quantitative assessment of HuR localization was done using our previously published method [10]. Fiji software was utilized for image analysis, and a minimal threshold was established in order to quantify the mean fluorescence intensity (FI). FI was quantified and normalized with the DAPI FI for the same region and expressed as arbitrary units (AU). Total lesion area was determined based on the ratio of GFAP-stained area (which is concentrated on the border of the injured region) to total spinal cord area. The total area of GFAP and myelin staining was measured using the freehand selection tool and threshold functions in Fiji. Three biological replicates were used for the Fiji analysis.

### Analysis of microglial morphology

During the early stages of activation, microglia can become hyper-ramified with increase branch complexity and soma enlargement [18, 19]. Fiji software (https://imagej.net/) was used to convert all photomicrographs to binary and skeletonized images using AnalyzeSkeleton plugin (https://imagej.net/plugins/analyze-skeleton/). To analyze the number of branches, the total branch length, and the endpoints per microglia brightness and contrast are changed by updating the Fiji lookup table, so pixel values are unchanged. This is followed by a despeckle step to remove salt-and-pepper noise. Photomicrographs then converted to binary using threshold tool followed by skeletonizing the image using the toolbar Skeletonize and AnalyzeSkeleton (2D/3D) to collect data on the branch length. Branch length per cell was calculated by dividing summed branch length by the number of microglia somas in the corresponding photomicrographs. Data was obtained from 5007 microglial cells from vehicle control (n=3) and 4538 microglial cells from SRI (n=3). The soma area was calculated using the Fiji multipoint area selection tool.

### Open field test

Motor function of the hind limbs was scored at 1, 2 and 3 weeks after SCI using the open-field basso mouse scale (BMS) in 10 min observation periods. BMS is a reliable measure and has been widely used to evaluate motor functions after SCI in mice [20]. Briefly, each mouse was separately placed in an open field (60 cm x 120 cm) and patterns of limb movement such as ankle movement, plantar placing and stepping of paw, paw positions and trunk instability were recorded over a 10 min time interval by a ceiling camera and analyzed by a computerized tracking system EthoVision XT sofware (Noldus, USA). Prior to the testing day, baseline measurements were obtained. Videos were carefully inspected by two reviewers (MAH and HAZ) blinded to the experimental conditions. Hindlimb joint movement, coordination, and weight support were evaluated using a rating scale from 0 points (no movement of any kind) to 9 points (normal locomotion). EthoVision software was used to analyze recorded videos to automatically calculate total distance traveled, mobility and velocity by each mouse within the arena.

### Beam walking

The apparatus consists of a round horizontal beam of 100 cm long and 1.2 cm in diameter and was elevated 50 cm above the ground. A hollow black escape box was attached to one end of the beam. One fluorescent lamp (60W) was used to illuminate the beam from the start side. One week prior to SCI, mice were thoroughly trained to traverse the beam from start point to end point (black box). Baseline measurements were taken one day before the SCI. On the trial days after SCI mice were placed at the start point and were allowed to walk for 2 min for 3 trials. The test was recorded by video camera. A scoring was given to the mice as described elsewhere [21].

Briefly, the rating system employs values such as mouse retention, forward motion, and goal-achieving on a beam. Mice received a high score for effectively using its hindlimbs to move around the beam. In case of absence of hindlimb movements, mice received no points. Scoring was done by an observer blinded to the identity of the mouse using recorded video weeks 1, 2, and 3 after SCI. A mean value was calculated from the three trials for each mouse.

### Rotarod

Motor coordination was assessed in weeks 1, 2, and 3 post SCI on an accelerating rotarod (San Diego instruments CA, USA). Mice were trained one week before SCI by placing them on the revolving rotor (4 rpm) and allowing them to acclimate for 2 mins. Baseline measures were acquired one day prior to the contusion injury. On the day of testing day, the rod accelerated from 4 to 60 rpm over 2 mins and the latency to fall off was measured in 2 trials on each testing day, allowing them 5 minutes of rest with in between each trial. The average time for the 2 trials was used for statistical analysis to calculate a single score for each mouse.

### Non-evoked Pain testing

Mice were placed into tabletop plexiglas cubicles (12 x 8 x 5.5 cm) with perforated metal floors and clear walls. High-resolution (1920×1080) cameras (Sony Handycam model HDR-CX100) were set up in front and behind the cubicles as described elsewhere [22]. After 15 min of habituation, cameras were turned on to record for 15 mins. Recordings were taken prior to surgery (baseline, BL) and at days 2 and 9 post SCI. Still images with clear visuals of the face were taken once in each 2-minute period for each recording (8 images total) by an experimenter blinded to the group assignment. Images were added to a PowerPoint file, randomized, and scored by two experimenters blinded to the experimental conditions. Scores for action units were averaged within each recording session. Action unit scores were averaged to calculate a mouse grimace score (MGS).

### Statistics

All statistical analyses were performed by Graphpad (GraphPad Software Inc.). The mean values are shown on graphs along with error bars. Error bars indicate the standard deviation and standard error as indicated. An unpaired Welch’s t test was used for HuR localization, qPCR, western blot, FI, and microglia counting. Two-way ANOVA with Tukey post hoc test was used for CCR2. Data from all behavioral testing were analyzed using an unpaired Welch’s t test. A p-value less than 0.05 was considered statistically significant.

## Results

### SRI-42127 treatment improves functional recovery after SCI

Wild-type female mice were subjected to a mid-thoracic contusion injury and then treated with SRI-42127 every 6 h for 5 days, starting 1 h after injury. To assess and quantitate motor activity over a longer period of time, we used a computerized tracking system in an open field **(Fig. 1A-E**). We first assessed locomotor function using the Basso mouse scale (BMS) at weeks 1, 2, and 3 post SCI **(Fig. 1A)**. There was a highly significant increase in BMS scores for SRI-treated mice versus vehicle beginning at week 1 (4.2 ± 1.2 versus 0.2 ± 0.1; *P* < 0.0001). The separation of BMS scores persisted and remained significant in weeks 2 and 3 with both groups showing some recovery by week 3. Heat maps of activity within the field were generated (example shown in **Fig. 1B**) from which distance moved, mobility and velocity were calculated **(Fig. 1C, D, E)**. At week 1, there was more than a two-fold increase in distance moved (3900 ± 200 versus 1800 ± 100 cm; *P* < 0.0001), mobility (6.5 ± 0.7 versus 4.0 ± 0.5 percent; *P* = 0.0029), and velocity (6.8 ± 1.0 versus 3.0 ± 0.5 cm/sec; *P* < 0.0001). The differences between SRI-and vehicle-treated groups remained significant throughout the recovery period for each parameter with both groups showing improvement by week 3. We next assessed the animals with a novel beam walking test which quantitates locomotor recovery using a scale of 0 to 7 **(Fig. 1F)** [21]. This test provides a more comprehensive assessment of beam walking by measuring other functional components besides latency to reach the other side. In both test groups, pre-injury baseline (BL) scores were 7. At week 1, there was a significant increase in score in the SRI-treated group compared to vehicle (3.3 ± 1.2 versus 0.6 ± 0.2; *P* < 0.0001). This difference persisted throughout the three-week observation period, with both groups showing modest improvement by week 3 (5.0 ± 1.2 versus 2.0 ± 0.5; *P* < 0.0001). Next, we assessed rotarod performance **(Fig. 1G)** and found that SRI-treated mice had a ∼5-fold increase in mean latency to fall at week 1 compared to vehicle control (6.8 ± 3.1 versus 1.3 ± 1; *P* = 0.0012). This difference remained significant throughout the recovery period with both groups showing increases in latency by week 3. To exclude the possibility of a drug effect on motor function, independent of SCI, we performed open field testing, BMS, and beam walking on sham-injured mice given SRI-42127 and observed no difference compared to the vehicle group over the 3-week testing period **(Supplementary Fig. 1**). Since pain is a common and debilitating sequelae of SCI, we assessed non-evoked pain using the mouse grimace scale (MGS) **(Fig. 1H)**. At baseline, prior to SCI, the groups had similar MGS scores (0.31 ± 0.02 versus 0.29 ± 0.04). Following SCI, mice treated with SRI-42127 had significantly lower MGS scores when compared to vehicle-treated mice (day 2: 1.56 ± 0.11 versus 1.02 ± 0.09, p < 0.01; day 9:0.83 ± 0.08 versus 0.46 ± 0.05, p < 0.01). Taken together, SRI-42127 treatment after SCI significantly attenuated motor function loss and pain.

**Figure 1:**
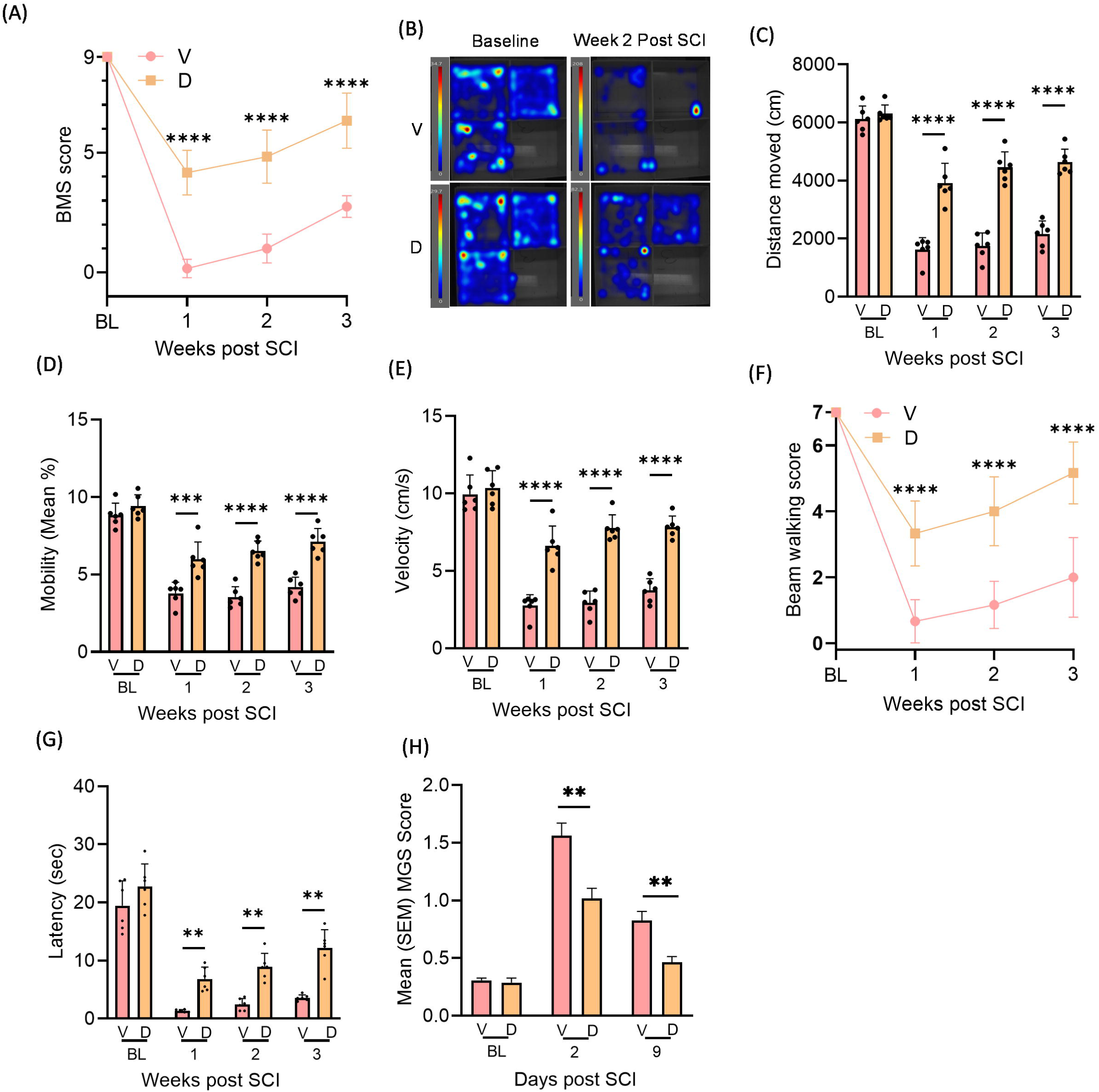
SRI-42127 attenuates motor function loss and pain after SCI. Following SCI, SRI-42127 (10mg/kg) or vehicle was administered IP at 6 h intervals for 5 days starting 1 h after injury. **(A)** BMS scores for vehicle control and SRI-treated mice were measured at 1, 2 and 3 weeks after SCI. **(B)** Motor function post SCI was assessed by open field testing. A representative heat map of activity at 2 weeks post SCI is shown. Different parameters of motor activity were measured including **(C)** distance moved, **(D)** mobility, and **(E)** velocity. **(F)** Beam walking scores of vehicle and SRI-treated mice were obtained at 1, 2, 3 weeks post SCI. **(G)** Rotarod testing was performed at 1-, 2- and 3-weeks post SCI and latency to fall was measured. **(H)** Non-evoked pain was assessed by mouse grimace scores (MGS) which were measured at day 2 and 9 post SCI. Baseline testing of both test groups, prior to SCI, had similar MGS scores. Evaluations for open field BMS, beam walking, and non-evoked pain were made by two observers blinded to the status of the animal. Error bars represent means ± SD of 6 mice/group. P values: * < 0.05, ** < 0.01, *** < 0.001, **** < 0.0001.

### SRI-42127 mitigates histopathology after SCI

At 3 weeks post SCI, spinal cords were harvested and tissue sections at the epicenter and adjacent rostral and caudal levels to assess for histopathological changes. To identify the glial scar border, we immunostained sections with GFAP **(Fig. 2A)**. Using these borders, a lesion size was calculated and found to be reduced by 60% in SRI-treated mice versus vehicle (P = 0.003) **(Fig. 2B)**. Inspection of rostral and caudal levels revealed no evidence of glial scar **(Supplementary Fig. 2A)**. To assess for neuronal loss, sections were immunostained with NeuN **(Fig. 2C)**. In SRI-treated mice, there was a 7-fold increase in NeuN positive cells in the epicenter (**Fig. 2D**; P= 0.003). Overall NeuN fluorescence intensity (FI) was significantly higher in SRI-treated mice **(Fig. 2E)**. In rostral and caudal levels, there were no differences in NeuN FI in SRI- and vehicle-treated mice **(Supplementary Fig. 2B**). To assess white matter sparing at 3 weeks in SRI-treated mice compared to vehicle controls (**Fig 2F, G**; *P* = 0.003). In rostral and caudal post SCI, spinal cord sections were stained with fluoromyelin. There was a 4.5-fold increase in FI levels, there was also a significant 40-50% increase in FI in SRI-treated mice (**Supplementary Fig. 2C**).

**Figure 2:**
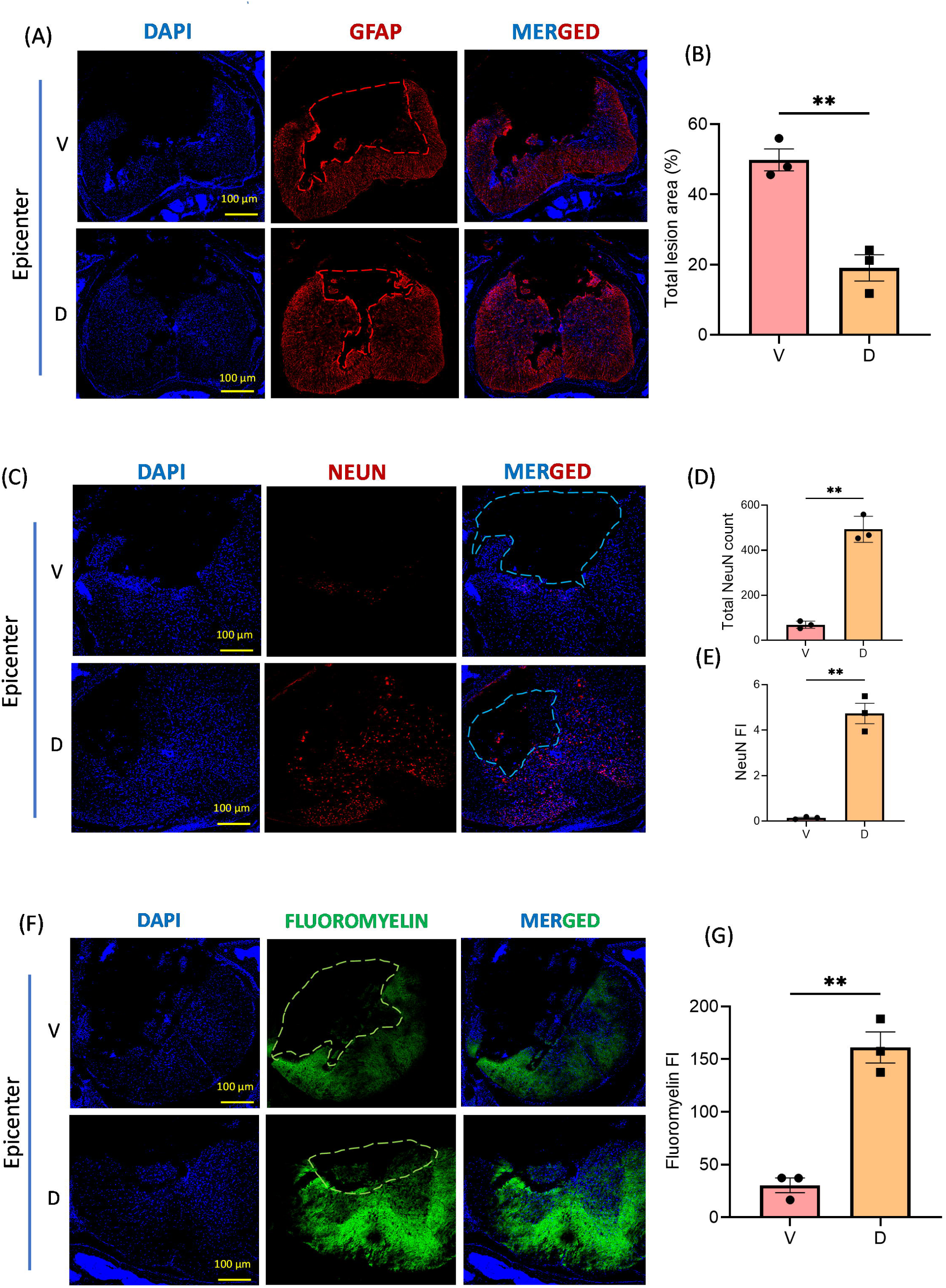
SRI-42127 treatment reduces lesion size and loss of neurons after SCI while increasing myelin sparing. **(A)** Representative GFAP-immunostained sections from the epicenter (3 weeks post SCI) showing reduced lesion size in an SRI-treated mouse compared to vehicle control. **(B)** Total lesion area was quantified in 3 vehicle and 3 SRI-treated mice. **(C)** Representative NeuN-immunostained sections from the epicenter showing an increase in NeuN+ cells in the SRI-treated mouse. **(D)** NeuN+ cells were quantified at the epicenter in 3 vehicle and 3 SRI-treated mice at 3 weeks post-SCI. **(E)** NeuN fluorescence intensity (FI) was quantified at the epicenter. **(F)** Representative fluoromyelin-stained sections from the epicenter showing increased myelin staining in SRI-treated mouse compared to vehicle control. **(G)** Fluoromyelin staining at the epicenter was quantified in 3 vehicle and 3 SRI-treated mice. Dashed lines delineate the boundary of injured spinal cord. Bars represent the mean ± SD of 3 mice/group. P < **0.01. Scale bars 100 μm.

### SRI-42127 blocks nucleocytoplasmic translocation of HuR in microglia and attenuates microglial activation

A hallmark of HuR activation is its translocation from nucleus to the cytoplasm where it stabilizes and promotes the translational efficiency of pro-inflammatory mediators in microglial cells [4, 10]. Here we assessed HuR localization by immunohistochemistry in Iba1+ cells at 8 h post SCI. At the epicenter, there was prominent HuR translocation in vehicle-treated mice, as indicated by a merged signal with cytoplasmic Iba1 (**Fig. 3A**). This shift was not seen in uninjured rostral or caudal levels (**Supplementary Fig. 3**). In SRI-treated mice, this merged signal was not observed (**Fig. 3B**). A nuclear/cytoplasmic ratio of HuR FI was calculated and found to be more than 2-fold higher in SRI-treated mice (*P* = 0.008) consistent with nuclear retention of HuR (**Fig. 3C**). To assess microglial activation, we measured Iba1 FI for individual cells in the epicenter and found an overall reduction consistent with reduced activation (**Fig. 3D**). This was underscored by a reduction in total Iba1 FI at the epicenter in SRI-treated mice (**Fig. 3E**) since the overall Iba1+ cell count was unchanged (**Fig. 3F**). Iba1 FI in rostral and caudal segments showed no differences between SRI-treated and vehicle control mice (**Fig. 3E**). Interestingly, we did not see translocation of HuR in astrocytes in vehicle-treated mice at this early timepoint but did at 24 h (**Supplementary Fig. 4A versus 4B**). In summary, these findings indicate prominent HuR cytoplasmic translocation in microglia (and astrocytes) after SCI which is suppressed with SRI-42127 treatment.

**Figure 3:**
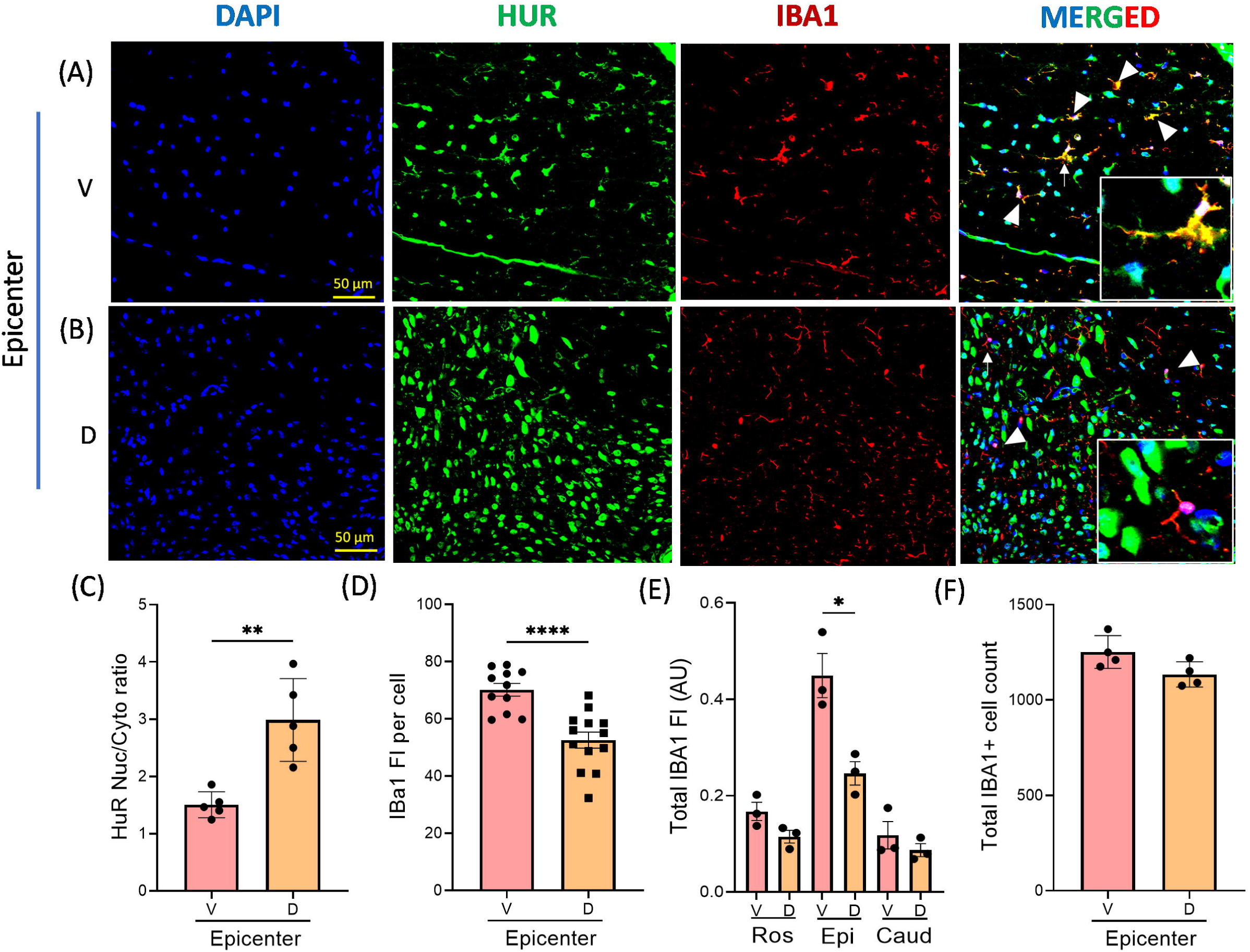
SRI-42127 inhibits nucleocytoplasmic translocation of HuR in microglia and attenuates microglial activation. Eight hours after SCI, sections from the epicenter were immunostained with antibodies as shown. **(A)** Cytoplasmic translocation of HuR is observed in microglia at the epicenter from a vehicle treated mouse as indicated by a merged yellow signal of HuR and Iba1 (arrow heads). A higher power view of a microglial cell (white arrow) is shown in the insert. **(B)** SRI-42127 treatment blocked HuR translocation from the nucleus as indicated by a merged signal of HuR, Iba1 and nuclear DAPI as well as an absence of a merged yellow signal. A higher power view of a microglial cell (white arrow) is shown in the insert. **(C)** Microglial HuR localization was quantified in 5 cells from 3 biological replicates as described in the Methods and expressed as a nuclear/cytoplasmic ratio. **(D)** Quantification of Iba1 fluorescence intensity (FI) in 11-13 cells from 3 biological replicates. **(E)** Total Iba1 FI was quantified at the epicenter, rostral, and caudal levels. **(F)** Total Iba1+ cell counts at the epicenter were quantified in 3 biological replicates. P * < .05, P < **0.01, *** < 0.001, **** < .0001. Scale bars 50 μm, 10 μm (inserts).

### SRI-42127 attenuates activation-associated microglia morphological changes

As another measure of activation, we assessed morphological changes of microglia at the epicenter 8 h after SCI. Using Fiji software, a skeleton analysis of microglia was done as previously described [23] (**Fig. 4A**). We observed a decrease in branch length (*P* = 0.009), number of branches (*P* < 0.0001), and number of end points per microglia (*P* < 0.0001) in drug-treated mice, consistent with reduced early-stage activation [24]. In addition to hyper-ramification, we measured soma size as an indicator of reactive microglia and found a significant reduction in SRI-treated mice (**Fig. 4B**; *P* < 0.0001). Taken together with results in Fig. 3, SRI-42127 attenuates microglial activation after SCI.

**Figure 4:**
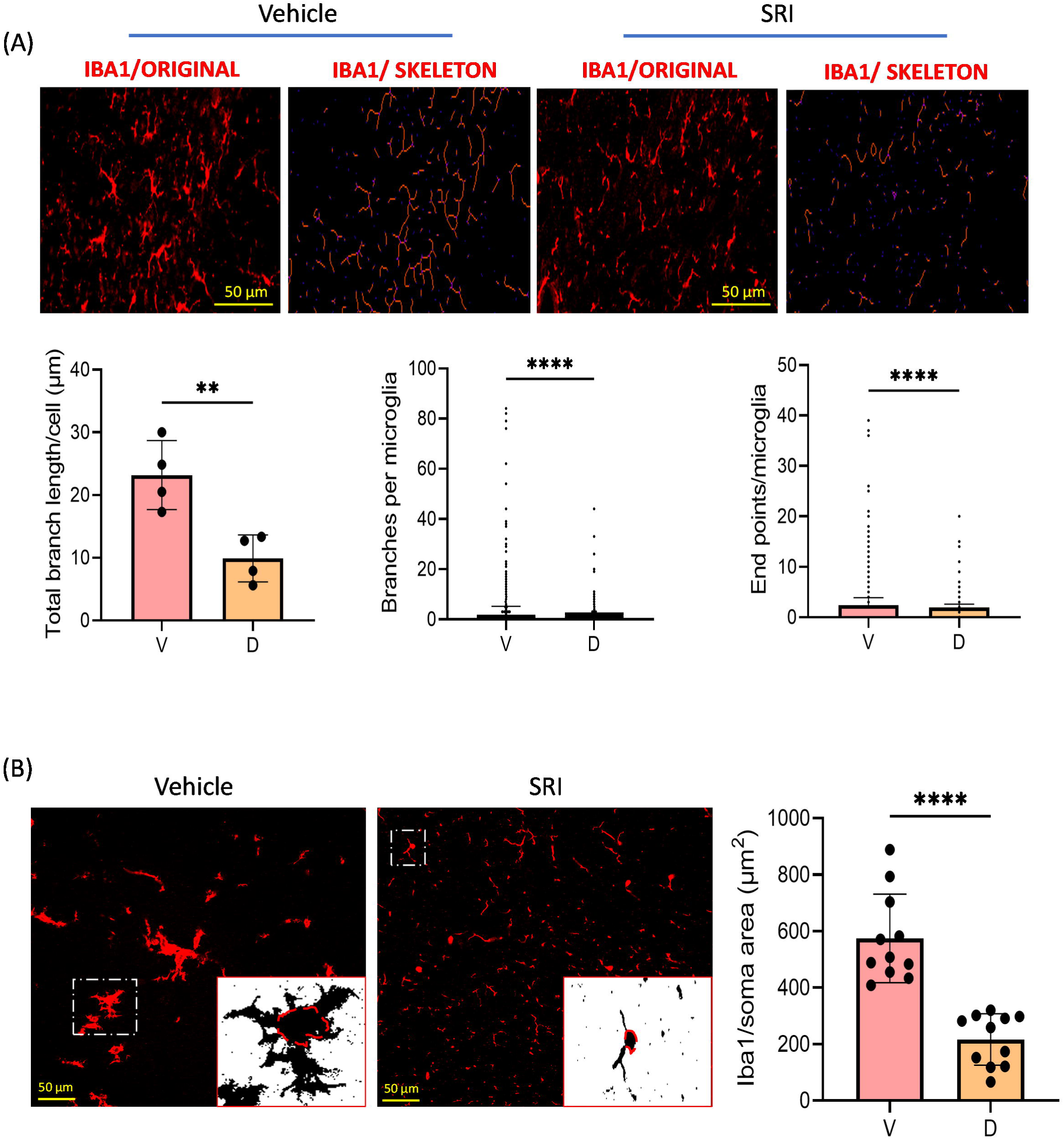
SRI-42127 attenuates activation-associated changes in microglial morphology after SCI. **(A)** Eight hours after SCI, tissue sections from the epicenter were immunostained with Iba1 antibodies and digitally converted to Skeletonized images using Fiji software. Representative images are shown. Morphological features were quantified including total microglia branch length, number of branches per microglial cell, and number of branch endpoints per microglial cell. Data were generated from 5007 microglial cells from vehicle control (n = 3 biological replicates) and 4538 microglial cells from SRI (n=3 biological replicates). **(B)** The soma of Iba1+ microglial cells were identified as shown in the representative images to the left. Binary images were generated by adjusting the Fiji threshold level (shown in inserts). Total area was quantified in 11 cells from 3 biological replicates as shown in the graph. Error bars represent SD. P *** < 0.001, **** < .0001.

### SRI-42127 suppresses induction of proinflammatory mediators at the epicenter of SCI

At 8 h post-SCI, spinal cords were harvested and divided into 3 segments: injury epicenter, rostral and caudal. RNA expression of proinflammatory mediators was assessed by qPCR. At the epicenter, there was a large induction of *IL-6*, *IL-1ꞵ*, *TNF-α*, *CCL2*, *MMP12*, *CXCL1*, *CXCL2*, and *iNOS* in the vehicle control compared to rostral and caudal segments (**Fig. 5A**). *COX2 and TGF-β1* mRNAs were not induced. The induction was most pronounced with *IL-6* (34-fold) and *CXCl2* (39-fold) while the others ranged from 5- to 15-fold. SRI-42127 treatment suppressed the expression of all pro-inflammatory mediators, most potently for *IL-6* (∼13-fold) and *IL-1β* (∼9-fold). Other inflammatory mediators were suppressed by 3- to 7-fold. Interestingly, there was SRI-induced suppression of some mRNAs at non-injured spinal cord segments, including *IL-1β*, *CCL2*, *MMP-12*, and *CXCL1*. Based on our prior work showing that HuR positively regulates *CCR2* in microglia [8], we assessed this mRNA after injury and found a nearly 3-fold attenuation at the epicenter. SRI-42127 suppressed *CCR2* mRNA at the uninjured levels as well. The fold-change was even larger as there was significantly higher expression of *CCR2* in non-injured levels for the vehicle control. Since COX-2 is heavily regulated by HuR at the translational efficiency level in addition to mRNA stability [25, 26], we assessed protein levels by western blot (**Fig. 5B**). We found a 2-fold increase in COX-2 protein levels in vehicle-treated mice compared to uninjured levels and this was attenuated by ∼4-fold in SRI-treated animals.

**Figure 5:**
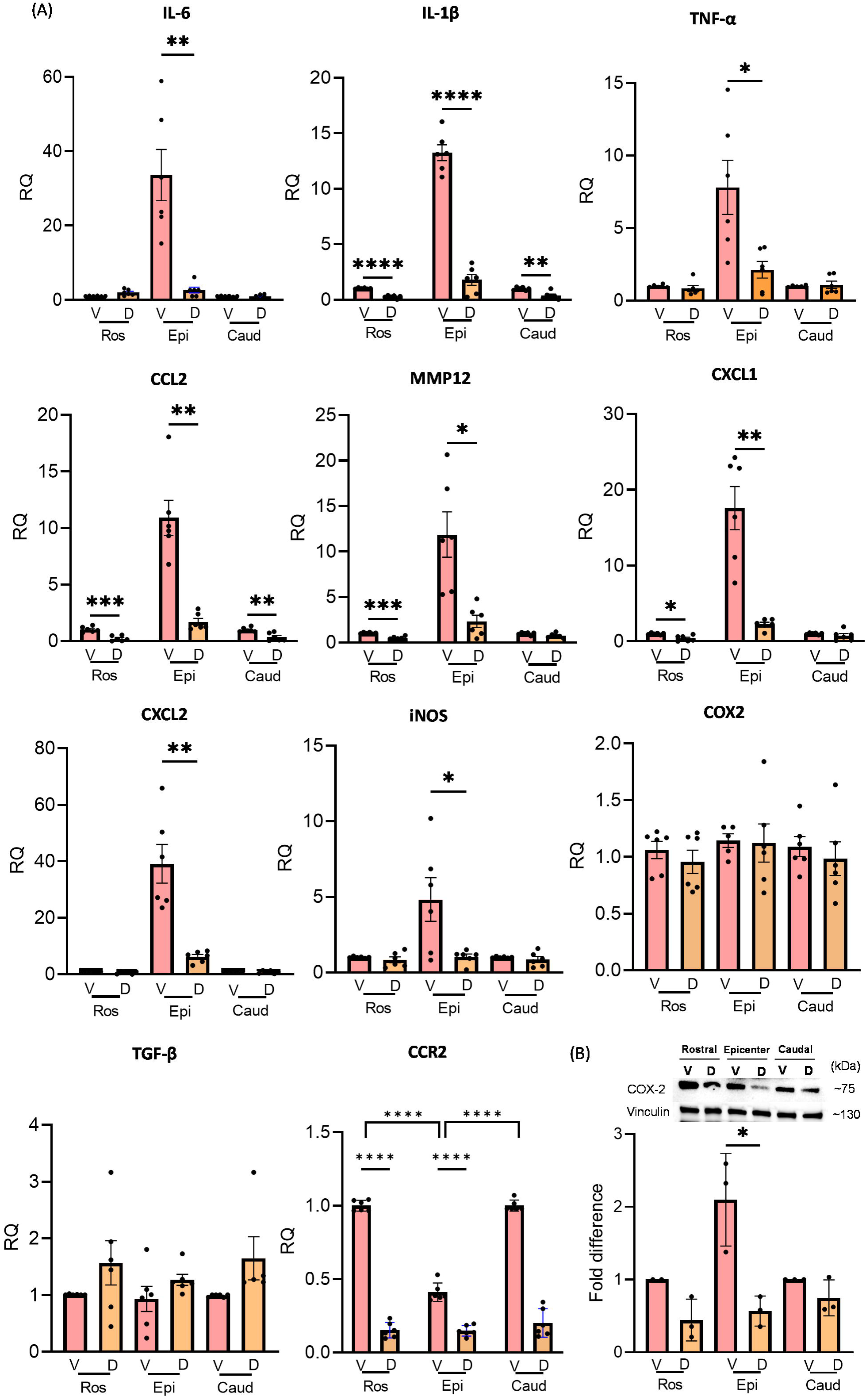
SRI-42127 attenuates pro-inflammatory cytokine and chemokine mRNA induction. **(A)** Spinal cords were harvested at 8 h post-SCI and divided into 3 segments: injury epicenter, rostral and caudal. RNA expression of proinflammatory cytokines and chemokines was assessed by qPCR in vehicle controls and SRI treated mice. All values were expressed relative to the vehicle control which was set at 1.0. GAPDH was used as the housekeeping control. P values: * < 0.05, ** < 0.01, *** < 0.001, **** < 0.0001. Error bars represent means ± SEM of 6 mice/group. **(B)** COX-2 western blot of protein lysate from epicenter, rostral and caudal spinal cord levels obtained 8 h after SCI. A representative blot is shown above. COX-2 levels were quantified by densitometry from three biological replicates using vinculin as a loading control. All values were expressed relative to the vehicle controls at rostral and caudal segments which were set at 1.0. Error bars represent the mean ± SD. P values: * < 0.05.

To determine the impact of SRI-42127 on inflammatory cytokine induction at a later timepoint, we assessed spinal cords after five days of SRI-42127 or vehicle treatment (**Supplementary Fig. 5**). There was a sustained suppression of pro-inflammatory mediators, most potently for *MMP-12* (∼90-fold) and *CXCl2* (∼30-fold). Others were suppressed by 3- to 9-fold. *TGF-β1*, on the other hand, was greater in vehicle-treated mice by ∼3-fold. No difference was seen with *IL-1β* while *CXCL1*, *iNOS*, and *COX-2* trended toward suppression but did not reach significance due to higher variability of mRNA levels. Taken together, treatment with SRI-42127 in the acute phase of SCI broadly suppressed the induction of pro-inflammatory mediators at the level of injury.

### SRI-42127 attenuates peripheral inflammatory responses after SCI

We next examined the effects of SRI-42127 on peripheral inflammatory responses at 8 h post SCI (**Fig. 6**). We first looked at circulating levels of key inflammatory mediators and found a large increase in IL-6 (5.8 vs. 6719 pg/ml) and CXCL1 (32.7 vs 3268) in vehicle-treated mice after SCI compared to sham-injured mice (**Fig. 6A**). CCL2 also increased by 3-fold (77.2 vs 226 pg/ml). With SRI-42127 treatment there was a significant attenuation of these inflammatory mediators: IL-6 (∼5-fold; *P* < 0.0001), CXCL1 (∼3.5-fold; *P* < 0.0001) and CCL2 (∼2.5-fold; *P* = 0.003). While there was some increase in IL-10, TNF-α, and IL1-β in vehicle-treated mice after SCI compared to sham-injured mice, SRI-42127 did not suppress this increase. To gain insight into potential sources for these cytokines, we looked at liver and spleen, two organs that drive the influx of inflammatory macrophages (**Fig. 6B**) [27, 28]. In liver tissue, we observed a significant induction of *CXCL1* (∼16-fold) and *IL-6* (9-fold) mRNA which was suppressed by ∼16-fold and 11-fold, respectively, with SRI-42127 treatment. For splenic tissue, there was an increase in *CCL2* (∼16-fold) and *IL-6* (∼10-fold) mRNA expression which was suppressed by ∼16-fold and ∼8-fold, respectively, with SRI-42127 treatment. For cytokines that were not elevated in serum, including IL-1β, TNF-α, and IL-10, there was only minimal induction of TNF-α in liver compared to sham injury **(Supplementary** Figure 6).

**Figure 6:**
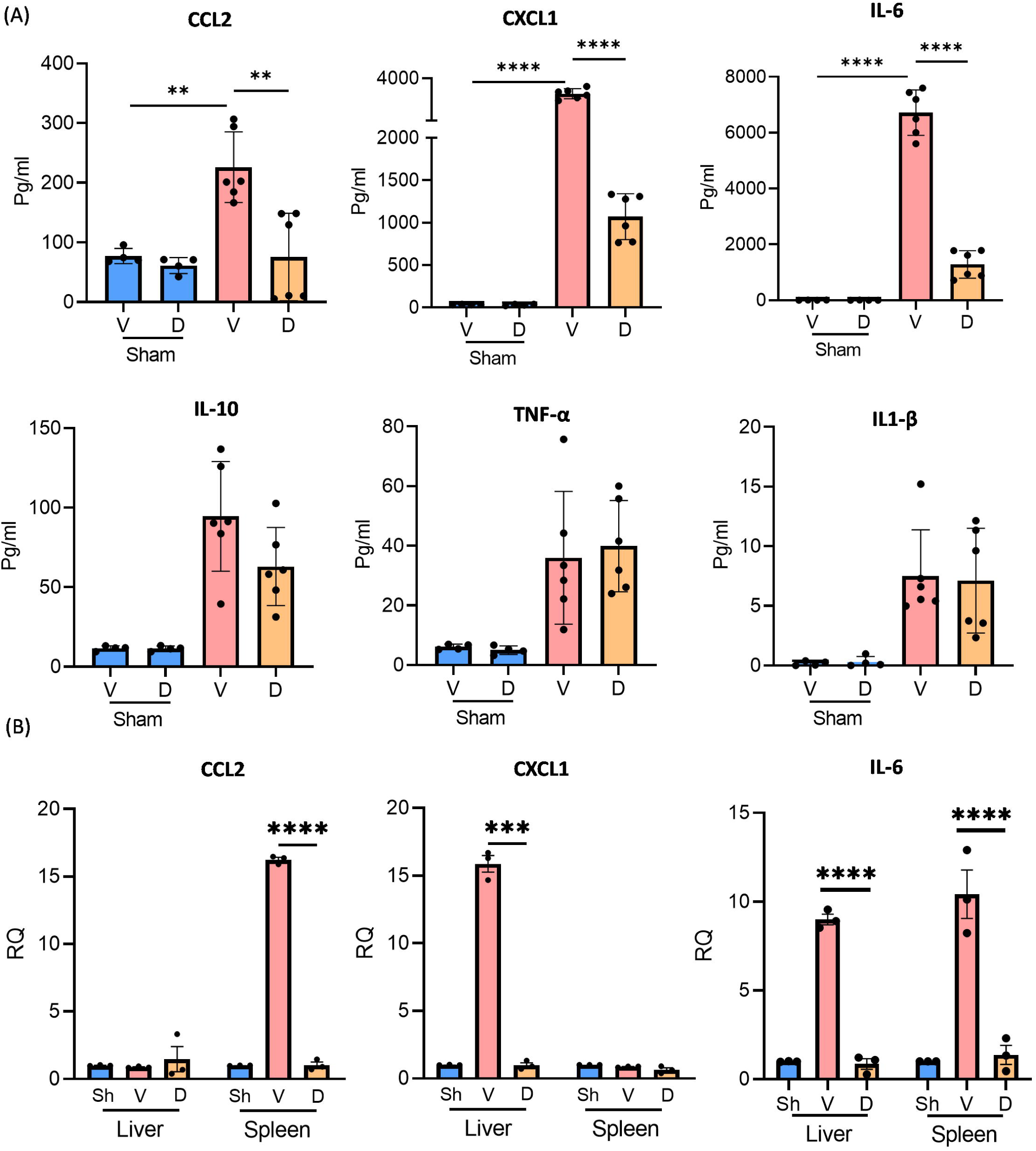
SRI-42127 suppresses expression of pro-inflammatory mediators peripherally. (A) ELISA was used to quantify cytokines in plasma samples obtained at 8 h post SCI or sham injury (Sh). **(B)** At the same time interval, RNA was harvested from liver and spleen and assessed by qPCR for *CXCL1*, *CCL2*, and *IL-6* mRNA expression. Data represent the mean ± SD of 6 mice/group in plasma and ± SD of 3 mice/group in liver and spleen. P values: ** < 0.01, *** < 0.001, **** < 0.0001.

## Discussion

A major challenge in therapeutically targeting the inflammatory cascade after SCI is the heterogeneity and complexity of signaling pathways driven by a broad range of pro-inflammatory mediators induced after injury [3, 29, 30]. These pathways are numerous, often overlapping, and orchestrated by many cell types at different stages of acute injury including microglia, macrophages, neutrophils, and astrocytes [30, 31]. HuR is a major RNA regulator of many pro-inflammatory mediators and is expressed in these cell types, making it a rational target for maximizing suppression of multiple components in the inflammatory cascade [3, 4, 7, 30, 32, 33]. In this report, we have shown that inhibiting HuR with the small molecule SRI-42127 potently suppresses pro-inflammatory responses in the spinal cord and periphery in the acute phase of SCI, resulting in a significant attenuation of motor function loss and pain. Histologic correlates for this therapeutic effect include sparing of neurons and whiter matter at the level of injury, reduced lesion size, and reduced microglial activation.

A deleterious effect of glial HuR in SCI was first reported by our group using a transgenic mouse model where its selective overexpression in astrocytes accentuated spinal cord edema, neuronal loss and neuroinflammatory responses after acute injury [9, 11]. In those reports, we described prominent nucleocytoplasmic translocation of both endogenous (and transgenic) HuR similar to what we observed in microglia and astrocytes in the current work. In glial cells, HuR is predominantly nuclear in localization, but upon activation by triggers such as LPS [10], hypoxia [10], glioblastoma [7, 32], and amyotrophic lateral sclerosis [8], translocates to the cytoplasm to promote mRNA stabilization and translational efficiency of pro-inflammatory mediators [4, 32]. HuR translocation was prominent in microglia/macrophages and astrocytes at the epicenter early of injury, but not rostrally or caudally, indicating a specificity of activation to the level of trauma. In astrocytes, HuR translocation occurred by 24 h post SCI, consistent with our previous reports [9, 11], but not at 8 h, suggesting that microglial HuR is activated earlier on in SCI. In both cell types, however, SRI-42127 attenuated cytoplasmic HuR which is consistent with its mechanism of blocking HuR dimerization [10, 13–15].

Inhibition of HuR with SRI-42127 led to a striking suppression of two cytokines, IL-6 and IL-1β, at the epicenter, both of which are rapidly induced within 1-3 hours after injury [29]. These cytokines are considered key initiators of the inflammatory cascade and directly produce toxic effects on neurons, oligodendrocytes, endothelial cells, leading to apoptosis and necrosis [30]. They are mainly produced by microglia and astrocytes activated upon exposure to damage associated molecular patterns (DAMPs) produced by the injury [4, 30]. This profound suppression was consistent with our prior studies indicating that SRI-42127 had the greatest suppressive effect on IL-1β and IL-6 induction in primary microglia and astrocytes *in cellulo* after LPS stimulation, and spinal cord microglia *in vivo* after spared nerve injury (SNI) [10, 16]. In the SNI model, SRI-42127 treatment significantly reversed allodynic pain in acute and chronic phases of the injury [16]. IL-6 and its signaling pathway are causally linked to neuropathic pain in SCI [34–36]. In prior reports, blocking the pathway with a neutralizing antibody to the IL-6 receptor or inhibiting the downstream transducer, JAK2, abrogated the development of pain and improved functional recovery. Thus, the early and marked suppression of IL-6 in the epicenter of SRI-treated mice may underlie the significant attenuation of spontaneous pain observed after SCI.

SRI-42127 suppressed other pro-inflammatory mediators in the initial 8 h post injury that further enhance the inflammatory cascade and toxic microenvironment via different pathways, including iNOS (via nitric oxide), COX-2 (via PGE_2_), and MMP-12 [3, 29, 30, 37, 38]. Interestingly COX-2 protein but not mRNA was suppressed with SRI-42127 treatment. This may be related to a loss of translational efficiency rather than mRNA stability (and decrease in mRNA levels) as both processes represent two distinct posttranscriptional levels of regulation governed by HuR [4]. We have shown previously that these mediators are regulated by HuR in glial cells in different inflammatory environments and suppressed either by SRI-42127, MS-444 (another chemical HuR inhibitor [15]) or with HuR silencing/deletion [7–10, 16, 39]. In addition to direct toxic effects, these inflammatory mediators also trigger indirect toxicity by promoting vascular permeability, edema, ischemia, and hemorrhage [1, 30].

In the acute phase of injury, the inflammatory cascade is further amplified by glial and macrophage production of chemokines, particularly CXCL1, CXCL2 and CCL2 which promote recruitment and activation of neutrophils, monocyte/macrophages, and other immune cells from the periphery [1, 30]. These chemokines were likewise suppressed by more than 6-fold with SRI-42127 treatment. In a compressive model of SCI, depletion of CCL2+ microglia significantly attenuated neuropathic pain [40]. CCL2-depeleted microglia switched from a pro- to anti-inflammatory phenotype and demonstrated reduced migration and invasion properties. CCL2 also can activate spinal microglia when injected intrathecally and trigger behavioral patterns consistent with neuropathic pain [41]. Interestingly, our prior work showed that HuR is a major positive regulator of CCR2, the main receptor for CCL2, in microglia and macrophages [8]. This is consistent with the >60% attenuation of *CCR2* mRNA levels at the epicenter with SRI-42127 treatment. *CCR2* knockout abrogates neuropathic pain after nerve injury which may tie into the attenuation of pain observed here [42, 43]. Of note, the fold-suppression of *CCR2* mRNA with drug treatment was even greater at rostral and caudal levels (∼6-fold) due to the higher expression of *CCR2* mRNA in vehicle controls at those levels. This difference is likely due to injury-related loss of neurons, astrocytes and oligodendrocytes, all of which express CCR2 [44].

Interestingly, *TGF-β1* was not induced at 8 h post injury or altered with SRI-42127 treatment but increased by ∼5-fold in vehicle-treated mice at 5 days. This timeframe is consistent with a prior report where TGF-β1 induction occurred more gradually (over days) after SCI [45]. In prior *in vivo* studies, we did not see suppression of TGF-β1 in brain or spinal cord tissues in SRI-treated mice [10, 16] and so the increase observed here may relate to enhanced glial scar signaling pathways due to a larger lesion size. TGF-β1 promotes astrocyte proliferation and glial scar formation and the 5-day timepoint represents the early stage of that process [46–48].

In SCI (and other forms of CNS trauma), there is a rapid peripheral response in organs such as liver and spleen that strongly influences the pro-inflammatory microenvironment in the injured cord by promoting mobilization and recruitment of inflammatory monocyte/macrophages and neutrophils [27, 28, 49–52]. In liver and spleen there is production and release of chemokines CXCL1 and CCL2 that drive this recruitment. Consistent with prior reports [50, 52], we observed a significant increase in circulating CXCL1 at the early timepoint of 8 hours post injury with liver tissue showing a ∼15-fold induction of *CXCL1* mRNA. SRI-42127 potently suppressed both hepatic mRNA induction and circulating levels of CXCL1. Although early induction of *CCL2* in liver has been observed with compression spinal cord injury models and later time points [51, 53], we did not see this at 8 hours in our model. On the other hand, circulating CCL2 levels were increased and spleen tissue showed a selective ∼15-fold induction of *CCL2* mRNA. A prior report identified spleen as the major source for monocyte-derived macrophages recruited to the injured spinal cord and pre-SCI splenectomy led to improved recovery [54].The induction of CCL2 after SCI was attributable to myeloid populations within the spleen that can express this chemokine [55], but it may serve as a signal for mobilizing and recruiting bone marrow derived monocytes that eventually infiltrate the injured spinal cord. Thus, the suppression of peripheral CCL2 by SRI-42127 may further reduce the inflammatory milieu in the injured spinal cord.

IL-6 was induced both in liver and splenic tissues in conjunction with a very large increase in circulating levels. In liver, IL-6 induction is a component of inflammation that has been observed in animal models of acute SCI [49]. Selective induction of IL-6 in liver and spleen versus TNF-α and IL-1β has been observed in other models of immune activation [56, 57] although the mechanism remains unclear. Our findings do suggest, however, that these organs are major contributors to circulating IL-6 in early SCI. In humans, elevated serum IL-6 has been detected in the acute phase of SCI, with some studies correlating levels with clinical severity of injury (see review [58]). The attenuation of systemic IL-6 by SRI-42127 may also have contributed to the reduction in pain after SCI as discussed above. Taken together, the suppression of peripheral inflammatory responses by SRI-42127 underscores the broad reach of HuR in cell types outside of the CNS [4, 59].

The effectiveness of SRI-42127 in blocking the inflammatory cascade is borne out by the significantly improved histological and behavioral outcomes. There was attenuation of microglial activation in the injured spinal cord, and reduced lesion size with sparing of neurons and white matter. Interestingly, we also observed a significant increase in myelin sparing in rostral and caudal levels in SRI-treated mice. Loss of myelin in these levels is likely related to Wallerian degeneration in long white matter tracts both rostral and caudal to injury, reflective of distal axonal degeneration following separation of axons from their cell body [47, 60].

In our prior work, we found that SRI-42127 penetrates the CNS to therapeutic concentrations within 10-15 minutes after systemic delivery [13] and rapidly suppresses neuroinflammatory responses [10]. These are favorable features for clinical translation in acute SCI where “time is spine” has emerged as a central theme in clinical management [6]. Because SRI-42127 can be administered systemically, it could be given in the field by emergency personnel shortly after injury. In our experimental paradigm, SRI-42127 was given 1 hour after SCI for a duration of 5 days.

Neuroinflammatory responses, including activated microglia and macrophages, persist in chronic phases when their role switches to a restorative one, promoting neural plasticity, angiogenesis and stem cell proliferation [3, 61, 62]. HuR regulates trophic factors that promote these restorative functions such as Hif-1α, VEGF, and GM-CSF, so prolonged treatment with SRI-42127 may be harmful [32, 59, 63–65]. Even IL-6 signaling may promote SCI recovery and neural regeneration [58]. However, the major deleterious pro-inflammatory cytokines regulated by HuR, approach pre-injury levels by 7 days, providing a time frame for intervention with HuR inhibitors. Further studies will be required to delineate the optimal duration for SRI-42127 treatment after SCI.

## Ethics approval and consent to participate

The animal study was reviewed and approved by the UAB Institutional Animal Care and Use Committee and were carried out in accordance with relevant guidelines and regulations of the National Research Council Guide for the Care and Use of Laboratory Animals.

## Consent for publication

Not applicable

## Availability of data and materials

The data that support the findings of this study are available on request from the corresponding author. The data are not publicly available due to privacy or ethical restrictions.

## Authors’ contributions

MAH, PHK, and CPC contributed to the conception and design of the study. MAH, RS, RES, AK, YS, AZH, KAS, and JD performed experiments and acquired data. AG, SAA, LBN, NF contributed to data analysis and interpretation. MAH and PHK prepared figures and drafted the manuscript. All authors read and approved the final manuscript.

## Acknowledgements

We wish to thank the UAB Animal Behavioral Assessment Core Facility for assistance in testing mice. We also thank Carlos A. Toro, PhD for helpful discussions regarding methodology.

## Abbreviations

ARE: AU-rich elements
BL: Baseline
BMS: Basso mouse scale
BSCB: Blood-spinal cord barrier
CCL2: C-C Motif chemokine ligand 2
CCR2: C-C chemokine receptor type 2
COX-2: Cyclooxygenase-2
CXCL: chemokine (C-X-C motif) ligand
D: Drug (SRI-42127)
DAMPs: Damage-associated molecular patterns
DAPI: 4’,6-diamidino-2-phenylindole
FI: Fluorescence intensity
GFAP: Glial fibrillary acidic protein
HuR: Human antigen R
Iba1: Ionized calcium binding adaptor molecule 1
IL-6: Interleukin 6
IL1-β: Interleukin-1 beta
iNOS: Inducible nitric oxide synthase
IP: intraperitoneal
LPS: Lipopolysaccharide
MGS: Mouse grimace scale
MMP: Matrix metalloproteinase
NeuN: Neuronal nuclei
PBS: Phosphate buffer saline
RBP: RNA binding protein
SCI: Spinal cord injury
Sh: Sham
SNI: Spared nerve injury
SRI: SRI-42127
TGF-β1: transforming growth factor beta-1
TNF-α: Tumor necrosis factor alpha
UTR: Untranslated region
V: Vehicle

## References

1. Alizadeh A, Dyck SM, Karimi-Abdolrezaee S: Traumatic Spinal Cord Injury: An Overview of Pathophysiology, Models and Acute Injury Mechanisms. Front Neurol 2019, 10.

2. Bennett J, J MD, Emmady PD: Spinal Cord Injuries. In StatPearls (Internet). Treasure Island (FL): StatPearls Publishing LLC.; 2024

3. Hellenbrand DJ, Quinn CM, Piper ZJ, Morehouse CN, Fixel JA, Hanna AS: Inflammation after spinal cord injury: a review of the critical timeline of signaling cues and cellular infiltration. J Neuroinflammation 2021, 18:284.

4. Guha A, Husain MA, Si Y, Nabors LB, Filippova N, Promer G, Smith R, King PH: RNA regulation of inflammatory responses in glia and its potential as a therapeutic target in central nervous system disorders. Glia 2023, 71:485–508.

5. Kumar H, Ropper AE, Lee SH, Han I: Propitious Therapeutic Modulators to Prevent Blood-Spinal Cord Barrier Disruption in Spinal Cord Injury. Mol Neurobiol 2017, 54:3578–3590.

6. Ahuja CS, Wilson JR, Nori S, Kotter MRN, Druschel C, Curt A, Fehlings MG: Traumatic spinal cord injury. Nature Reviews Disease Primers 2017, 3:17018.

7. Wang J, Leavenworth JW, Hjelmeland AB, Smith R, Patel N, Borg B, Si Y, King PH: Deletion of the RNA regulator HuR in tumor-associated microglia and macrophages stimulates anti-tumor immunity and attenuates glioma growth. Glia 2019, 67:2424–2439.

8. Matsye P, Zheng L, Si Y, Kim S, Luo W, Crossman DK, Bratcher PE, King PH: HuR promotes the molecular signature and phenotype of activated microglia: Implications for amyotrophic lateral sclerosis and other neurodegenerative diseases. Glia 2017, 65:945–963.

9. Kwan T, Floyd CL, Kim S, King PH: RNA Binding Protein Human Antigen R Is Translocated in Astrocytes following Spinal Cord Injury and Promotes the Inflammatory Response. J Neurotrauma 2017, 34:1249–1259.

10. Chellappan R, Guha A, Si Y, Kwan T, Nabors LB, Filippova N, Yang X, Myneni AS, Meesala S, Harms AS, King PH: SRI-42127, a novel small molecule inhibitor of the RNA regulator HuR, potently attenuates glial activation in a model of lipopolysaccharide-induced neuroinflammation. Glia 2022, 70:155–172.

11. Kwan T, Floyd CL, Patel J, Mohaimany-Aponte A, King PH: Astrocytic expression of the RNA regulator HuR accentuates spinal cord injury in the acute phase. Neurosci Lett 2017, 651:140–145.

12. Guo H, Du M, Yang Y, Lin X, Wang Y, Li H, Ren J, Xu W, Yan J, Wang N: Sp1 regulates the M1 polarization of microglia through the HuR/NF-κB axis after spinal cord injury. Neuroscience 2024.

13. Filippova N, Yang X, Ananthan S, Calano J, Pathak V, Bratton L, Vekariya RH, Zhang S, Ofori E, Hayward EN, et al: Targeting the HuR Oncogenic Role with a New Class of Cytoplasmic Dimerization Inhibitors. Cancer Res 2021, 81:2220–2233.

14. Filippova N, Yang X, Ananthan S, Sorochinsky A, Hackney JR, Gentry Z, Bae S, King P, Nabors LB: Hu antigen R (HuR) multimerization contributes to glioma disease progression. J Biol Chem 2017, 292:16999–17010.

15. Meisner NC, Hintersteiner M, Mueller K, Bauer R, Seifert JM, Naegeli HU, Ottl J, Oberer L, Guenat C, Moss S, et al: Identification and mechanistic characterization of low-molecular-weight inhibitors for HuR. Nat Chem Biol 2007, 3:508–515.

16. Sorge RE, Si Y, Norian LA, Guha A, Moore GE, Nabors LB, Filippova N, Yang X, Smith R, Chellappan R, King PH: Inhibition of the RNA Regulator HuR by SRI-42127 Attenuates Neuropathic Pain After Nerve Injury Through Suppression of Neuroinflammatory Responses. Neurotherapeutics 2022, 19:1649–1661.

17. Guida F, De Gregorio D, Palazzo E, Ricciardi F, Boccella S, Belardo C, Iannotta M, Infantino R, Formato F, Marabese I, et al: Behavioral, Biochemical and Electrophysiological Changes in Spared Nerve Injury Model of Neuropathic Pain. In Int J Mol Sci, vol. 21, 2020/05/15 edition; 2020.

18. Ziebell JM, Adelson PD, Lifshitz J: Microglia: dismantling and rebuilding circuits after acute neurological injury. Metab Brain Dis 2015, 30:393–400.

19. Streit WJ, Walter SA, Pennell NA: Reactive microgliosis. Prog Neurobiol 1999, 57:563–581.

20. Basso DM, Fisher LC, Anderson AJ, Jakeman LB, McTigue DM, Popovich PG: Basso Mouse Scale for locomotion detects differences in recovery after spinal cord injury in five common mouse strains. J Neurotrauma 2006, 23:635–659.

21. Ito S, Kakuta Y, Yoshida K, Shirota Y, Mieda T, Iizuka Y, Chikuda H, Iizuka H, Nakamura K: A simple scoring of beam walking performance after spinal cord injury in mice. PLoS One 2022, 17:e0272233.

22. Langford DJ, Bailey AL, Chanda ML, Clarke SE, Drummond TE, Echols S, Glick S, Ingrao J, Klassen-Ross T, Lacroix-Fralish ML, et al: Coding of facial expressions of pain in the laboratory mouse. Nat Methods 2010, 7:447–449.

23. Green TRF, Murphy SM, Rowe RK: Comparisons of quantitative approaches for assessing microglial morphology reveal inconsistencies, ecological fallacy, and a need for standardization. Sci Rep 2022, 12:18196.

24. Reddaway J, Richardson PE, Bevan RJ, Stoneman J, Palombo M: Microglial morphometric analysis: so many options, so little consistency. Front Neuroinform 2023, 17:1211188.

25. Dixon DA, Tolley ND, King PH, Nabors LB, McIntyre TM, Zimmerman GA, Prescott SM: Altered expression of the mRNA stability factor HuR promotes cyclooxygenase-2 expression in colon cancer cells. J Clin Invest 2001, 108:1657–1665.

26. Cok SJ, Morrison AR: The 3’-Untranslated Region of Murine Cyclooxygenase-2 Contains Multiple Regulatory Elements That Alter Message Stability and Translational Efficiency. J Biol Chem 2001, 276:23179–23185.

27. Anthony DC, Couch Y: The systemic response to CNS injury. Exp Neurol 2014, 258:105–111.

28. Noble BT, Brennan FH, Popovich PG: The spleen as a neuroimmune interface after spinal cord injury. Journal of Neuroimmunology 2018, 321:1–11.

29. Sterner RC, Sterner RM: Immune response following traumatic spinal cord injury: Pathophysiology and therapies. Front Immunol 2022, 13:1084101.

30. Anwar MA, Al Shehabi TS, Eid AH: Inflammogenesis of Secondary Spinal Cord Injury. Front Cell Neurosci 2016, 10:98.

31. Moraga I, Spangler J, Mendoza JL, Garcia KC: Multifarious determinants of cytokine receptor signaling specificity. Adv Immunol 2014, 121:1–39.

32. Guha A, Waris S, Nabors LB, Filippova N, Gorospe M, Kwan T, King PH: The versatile role of HuR in Glioblastoma and its potential as a therapeutic target for a multi-pronged attack. Adv Drug Deliv Rev 2022, 181:114082.

33. Bonomo I, Assoni G, La Pietra V, Canarutto G, Facen E, Donati G, Zucal C, Genovese S, Micaelli M, Pérez-Ràfols A, et al: HuR modulation with tanshinone mimics impairs LPS response in murine macrophages. Disease Models & Mechanisms 2023.

34. Guptarak J, Wanchoo S, Durham-Lee J, Wu Y, Zivadinovic D, Paulucci-Holthauzen A, Nesic O: Inhibition of IL-6 signaling: A novel therapeutic approach to treating spinal cord injury pain. Pain 2013, 154:1115–1128.

35. Murakami T, Kanchiku T, Suzuki H, Imajo Y, Yoshida Y, Nomura H, Cui D, Ishikawa T, Ikeda E, Taguchi T: Anti-interleukin-6 receptor antibody reduces neuropathic pain following spinal cord injury in mice. Exp Ther Med 2013, 6:1194–1198.

36. Lee JY, Park CS, Seo KJ, Kim IY, Han S, Youn I, Yune TY: IL-6/JAK2/STAT3 axis mediates neuropathic pain by regulating astrocyte and microglia activation after spinal cord injury. Exp Neurol 2023, 370:114576.

37. Ji H, Zhang Y, Chen C, Li H, He B, Yang T, Sun C, Hao H, Zhang X, Wang Y, et al: D-dopachrome tautomerase activates COX2/PGE(2) pathway of astrocytes to mediate inflammation following spinal cord injury. J Neuroinflammation 2021, 18:130.

38. Chelluboina B, Nalamolu KR, Klopfenstein JD, Pinson DM, Wang DZ, Vemuganti R, Veeravalli KK: MMP-12, a Promising Therapeutic Target for Neurological Diseases. Mol Neurobiol 2018, 55:1405–1409.

39. Wang J, Hjelmeland AB, Nabors LB, King PH: Anti-cancer effects of the HuR inhibitor, MS-444, in malignant glioma cells. Cancer Biol Ther 2019, 20:979–988.

40. Li Q, Yang Z, Wang K, Chen Z, Shen H: Suppression of microglial Ccl2 reduces neuropathic pain associated with chronic spinal compression. Front Immunol 2023, 14:1191188.

41. Thacker MA, Clark AK, Bishop T, Grist J, Yip PK, Moon LD, Thompson SW, Marchand F, McMahon SB: CCL2 is a key mediator of microglia activation in neuropathic pain states. Eur J Pain 2009, 13:263–272.

42. Ma M, Wei T, Boring L, Charo IF, Ransohoff RM, Jakeman LB: Monocyte recruitment and myelin removal are delayed following spinal cord injury in mice with CCR2 chemokine receptor deletion. Journal of Neuroscience Research 2002, 68:691–702.

43. Abbadie C, Lindia JA, Cumiskey AM, Peterson LB, Mudgett JS, Bayne EK, DeMartino JA, MacIntyre DE, Forrest MJ: Impaired neuropathic pain responses in mice lacking the chemokine receptor CCR2. Proc Natl Acad Sci U S A 2003, 100:7947–7952.

44. Banisadr G, Quéraud-Lesaux F, Boutterin MC, Pélaprat D, Zalc B, Rostène W, Haour F, Mélik Parsadaniantz S: Distribution, cellular localization and functional role of CCR2 chemokine receptors in adult rat brain. Journal of Neurochemistry 2002, 81:257–269.

45. Joko M, Osuka K, Usuda N, Atsuzawa K, Aoyama M, Takayasu M: Different modifications of phosphorylated Smad3C and Smad3L through TGF-beta after spinal cord injury in mice. Neurosci Lett 2013, 549:168–172.

46. Gong L, Gu Y, Han X, Luan C, Liu C, Wang X, Sun Y, Zheng M, Fang M, Yang S, et al: Spatiotemporal Dynamics of the Molecular Expression Pattern and Intercellular Interactions in the Glial Scar Response to Spinal Cord Injury. Neurosci Bull 2023, 39:213–244.

47. Shafqat A, Albalkhi I, Magableh HM, Saleh T, Alkattan K, Yaqinuddin A: Tackling the glial scar in spinal cord regeneration: new discoveries and future directions. Front Cell Neurosci 2023, 17:1180825.

48. Ma CW, Wang ZQ, Ran R, Liao HY, Lyu JY, Ren Y, Lei ZY, Zhang HH: TGF-β signaling pathway in spinal cord injury: Mechanisms and therapeutic potential. J Neurosci Res 2024, 102:e25255.

49. Goodus MT, McTigue DM: Hepatic dysfunction after spinal cord injury: A vicious cycle of central and peripheral pathology? Exp Neurol 2020, 325:113160.

50. Yates AG, Jogia T, Gillespie ER, Couch Y, Ruitenberg MJ, Anthony DC: Acute IL-1RA treatment suppresses the peripheral and central inflammatory response to spinal cord injury. Journal of Neuroinflammation 2021, 18:15.

51. Campbell SJ, Perry VH, Pitossi FJ, Butchart AG, Chertoff M, Waters S, Dempster R, Anthony DC: Central nervous system injury triggers hepatic CC and CXC chemokine expression that is associated with leukocyte mobilization and recruitment to both the central nervous system and the liver. Am J Pathol 2005, 166:1487–1497.

52. Campbell SJ, Hughes PM, Iredale JP, Wilcockson DC, Waters S, Docagne F, Perry VH, Anthony DC: CINC-1 is an acute-phase protein induced by focal brain injury causing leukocyte mobilization and liver injury. Faseb j 2003, 17:1168–1170.

53. D’Mello C, Le T, Swain MG: Cerebral microglia recruit monocytes into the brain in response to tumor necrosis factoralpha signaling during peripheral organ inflammation. J Neurosci 2009, 29:2089–2102.

54. Blomster LV, Brennan FH, Lao HW, Harle DW, Harvey AR, Ruitenberg MJ: Mobilisation of the splenic monocyte reservoir and peripheral CX3CR1 deficiency adversely affects recovery from spinal cord injury. Experimental Neurology 2013, 247:226–240.

55. Gschwandtner M, Derler R, Midwood KS: More Than Just Attractive: How CCL2 Influences Myeloid Cell Behavior Beyond Chemotaxis. Frontiers in Immunology 2019, 10.

56. Szot P, Franklin A, Figlewicz DP, Beuca TP, Bullock K, Hansen K, Banks WA, Raskind MA, Peskind ER: Multiple lipopolysaccharide (LPS) injections alter interleukin 6 (IL-6), IL-7, IL-10 and IL-6 and IL-7 receptor mRNA in CNS and spleen. Neuroscience 2017, 355:9–21.

57. Zhu XL, Pacheco ND, Dick EJ, Rollwagen FM: Differentially increased IL-6 mRNA expression in liver and spleen following injection of liposome-encapsulated haemoglobin. Cytokine 1999, 11:696–703.

58. Shipman H, Monsour M, Foley MM, Marbacher S, Croci DM, Bisson EF: Interleukin-6 in Spinal Cord Injury: Could Immunomodulation Replace Immunosuppression in the Management of Acute Traumatic Spinal Cord Injuries? J Neurol Surg A Cent Eur Neurosurg 2024.

59. Srikantan S, Gorospe M: HuR function in disease. Front Biosci (Landmark Ed) 2012, 17:189–205.

60. Cohen-Adad J, Leblond H, Delivet-Mongrain H, Martinez M, Benali H, Rossignol S: Wallerian degeneration after spinal cord lesions in cats detected with diffusion tensor imaging. Neuroimage 2011, 57:1068–1076.

61. Wu Y, Tang Z, Zhang J, Wang Y, Liu S: Restoration of spinal cord injury: From endogenous repairing process to cellular therapy. Front Cell Neurosci 2022, 16:1077441.

62. Freyermuth-Trujillo X, Segura-Uribe JJ, Salgado-Ceballos H, Orozco-Barrios CE, Coyoy-Salgado A: Inflammation: A Target for Treatment in Spinal Cord Injury. Cells 2022, 11.

63. Li Y, Han W, Wu Y, Zhou K, Zheng Z, Wang H, Xie L, Li R, Xu K, Liu Y, et al: Stabilization of Hypoxia Inducible Factor-1alpha by Dimethyloxalylglycine Promotes Recovery from Acute Spinal Cord Injury by Inhibiting Neural Apoptosis and Enhancing Axon Regeneration. J Neurotrauma 2019, 36:3394–3409.

64. Zamanian C, Kim G, Onyedimma C, Ghaith AK, Jarrah R, Graepel S, Moinuddin FM, Bydon M: A review of vascular endothelial growth factor and its potential to improve functional outcomes following spinal cord injury. Spinal Cord 2023, 61:231–237.

65. Tao JW, Fan X, Zhou JY, Huo LY, Mo YJ, Bai HZ, Zhao Y, Ren JP, Mu XH, Xu L: Granulocyte colony-stimulating factor effects on neurological and motor function in animals with spinal cord injury: a systematic review and meta-analysis. Front Neurosci 2023, 17:1168764.

