## Supplementary figures and images for "Inhibition of the RNA Regulator HuR mitigates spinal cord injury by potently suppressing post-injury neuroinflammation"

### Supplementary Fig. 1

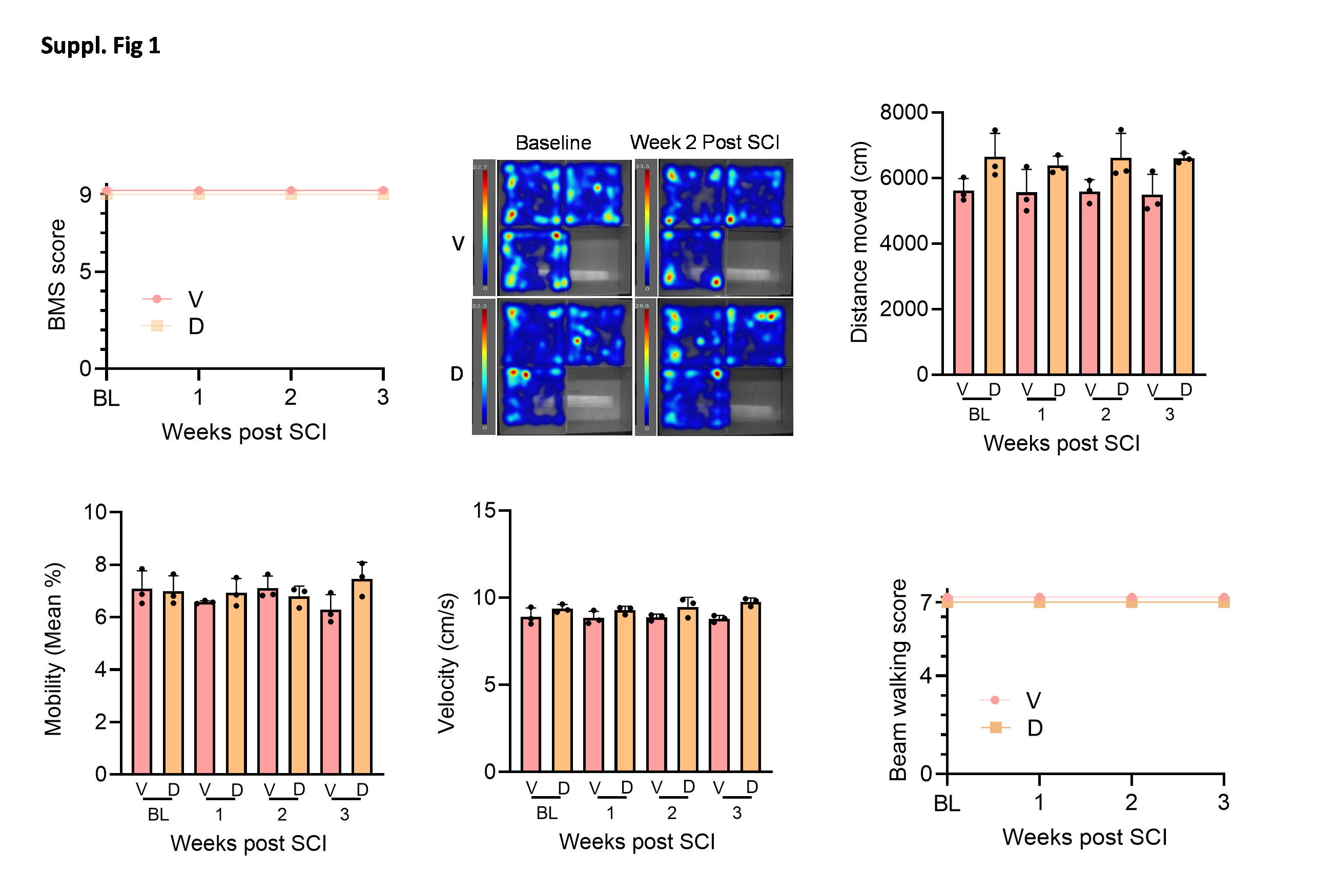

### Supplementary Fig. 2

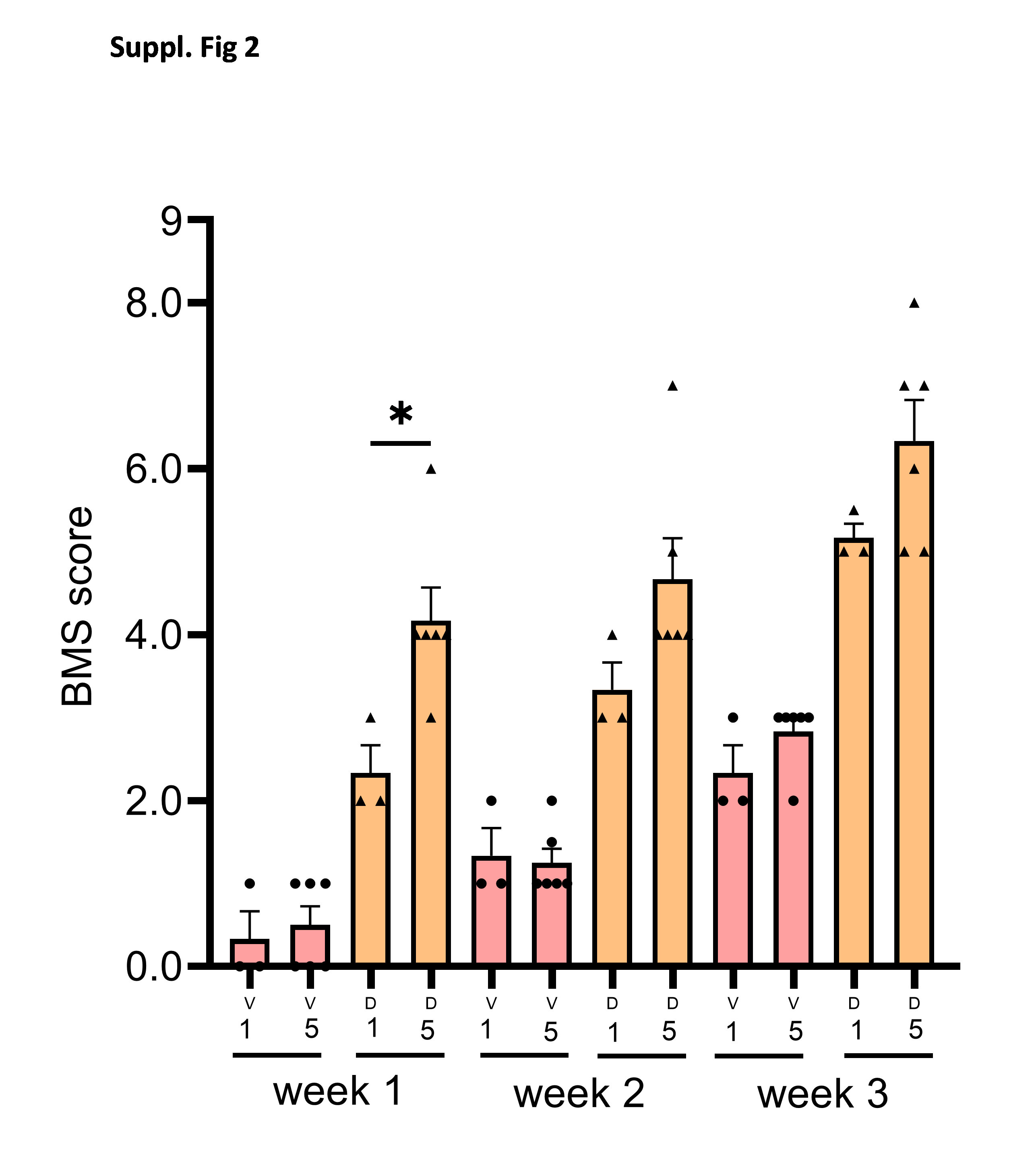

### Supplementary Fig. 3

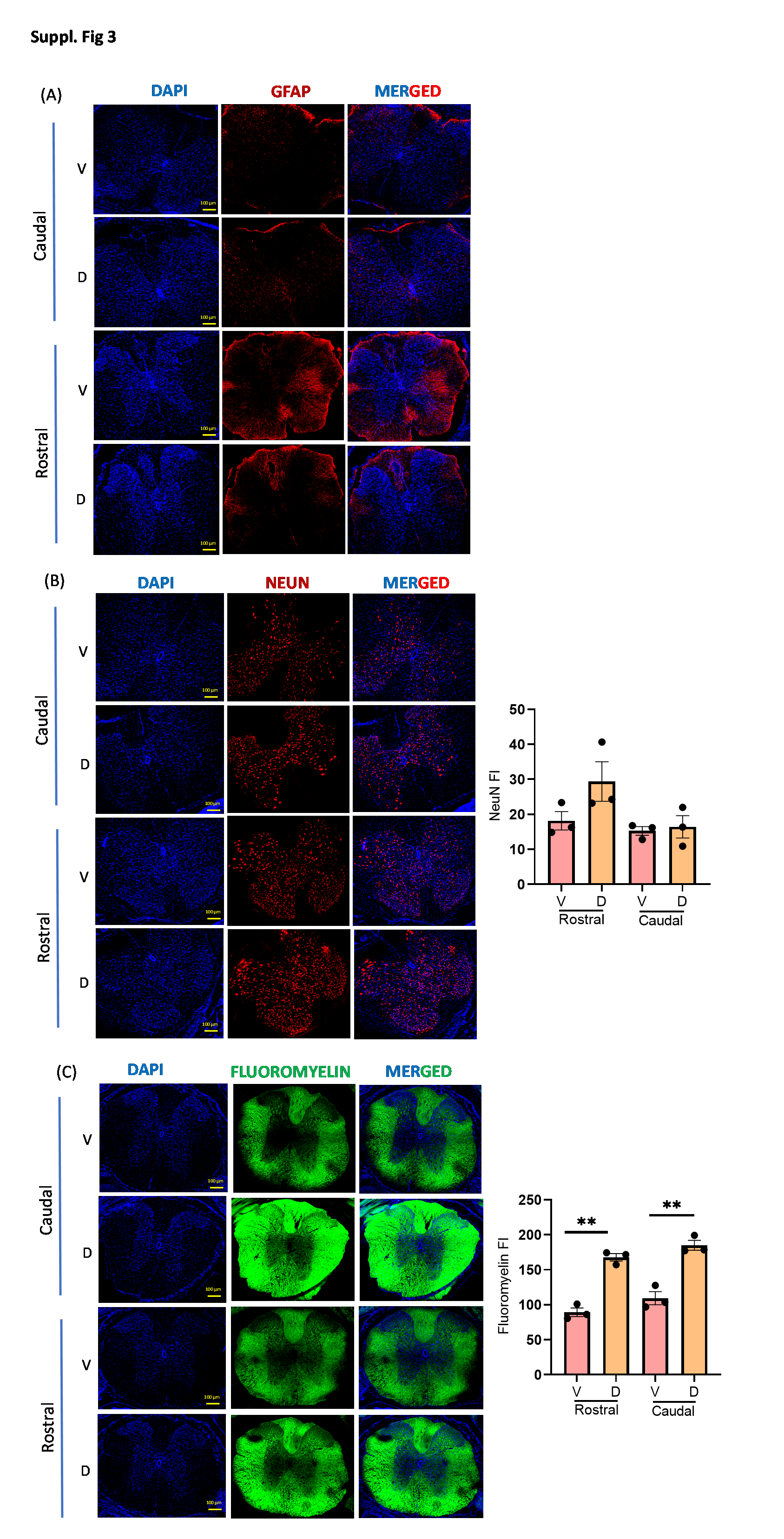

### Supplementary Fig. 4

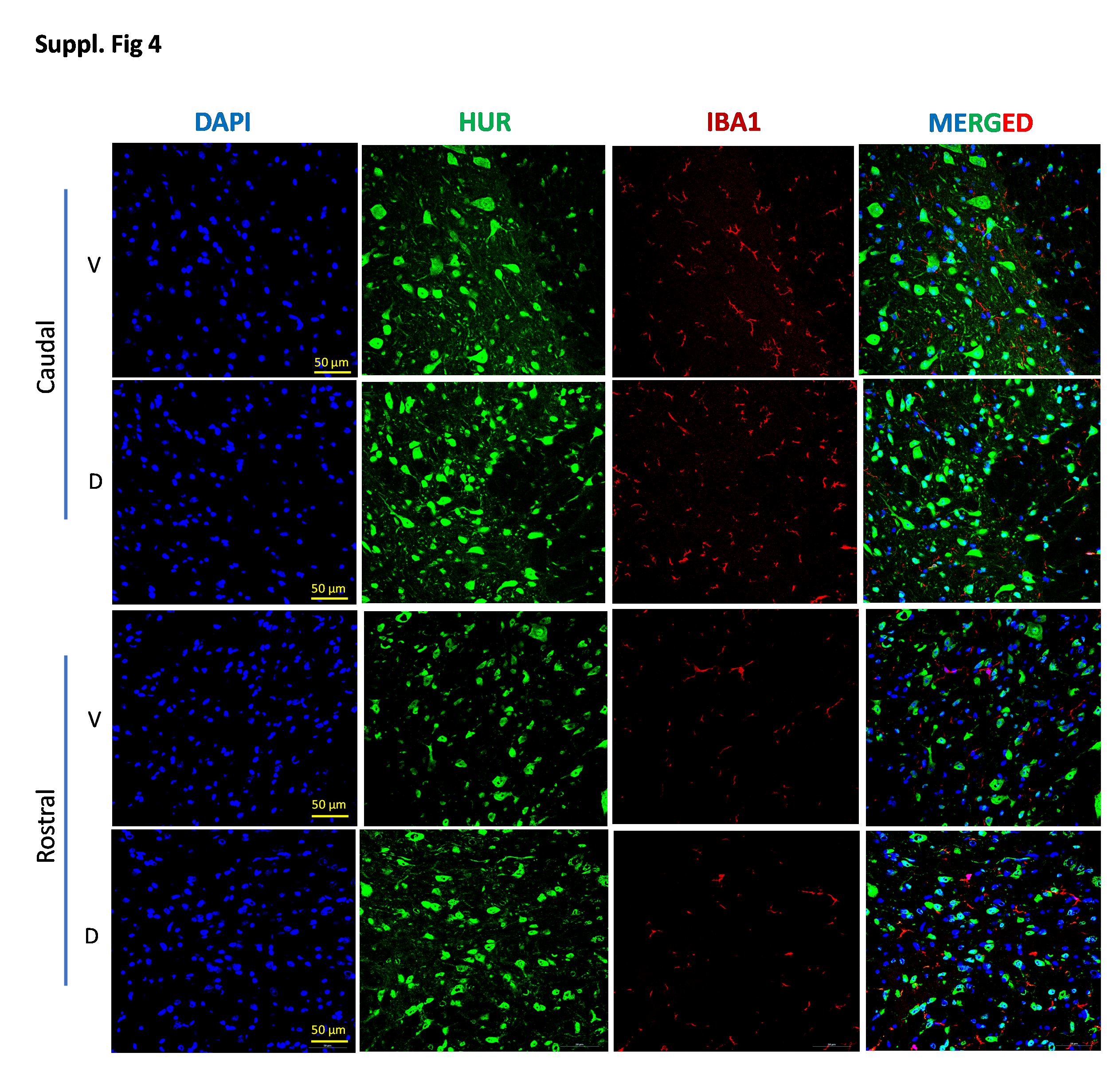

### Supplementary Fig. 5

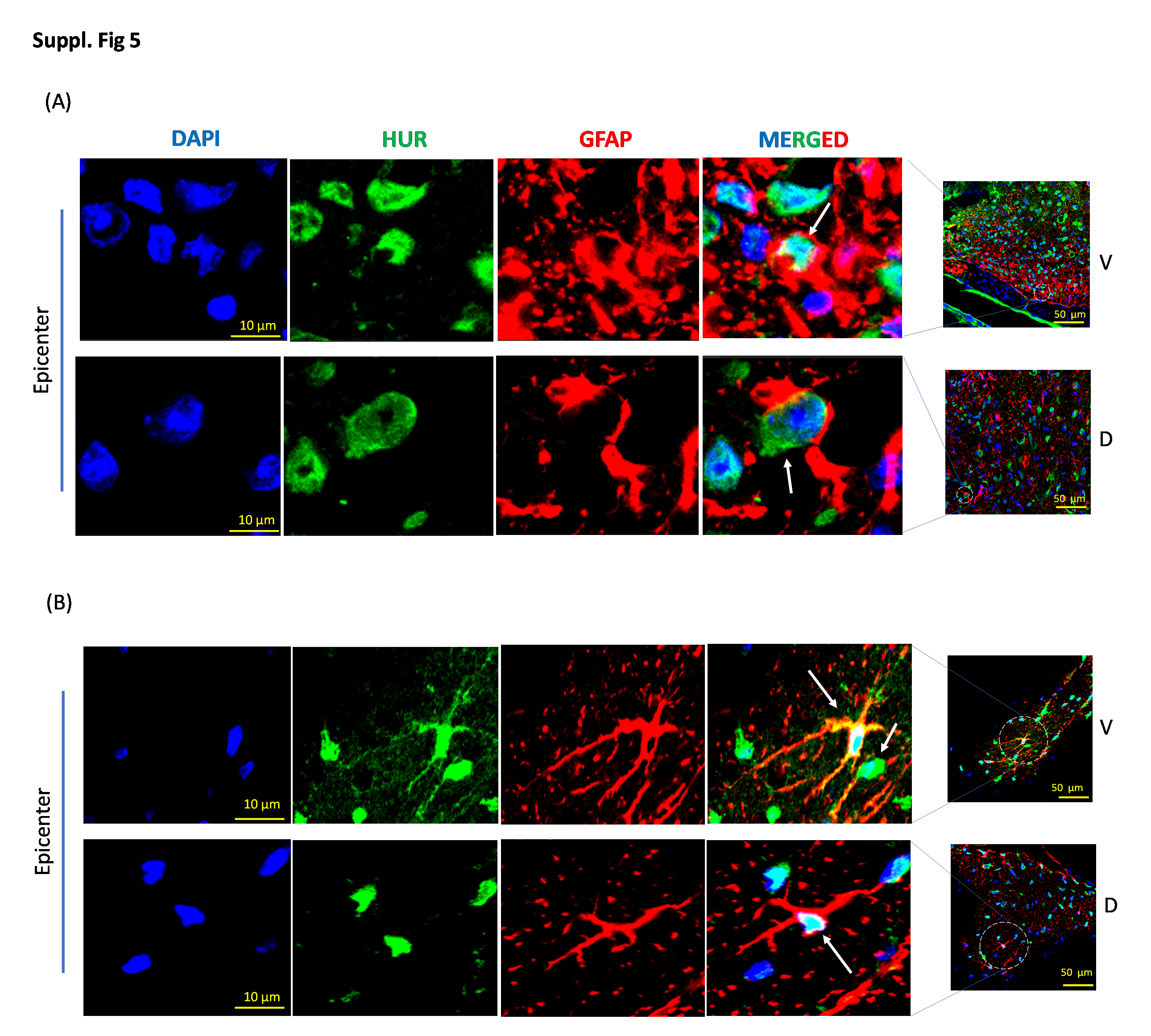

### Supplementary Fig. 6

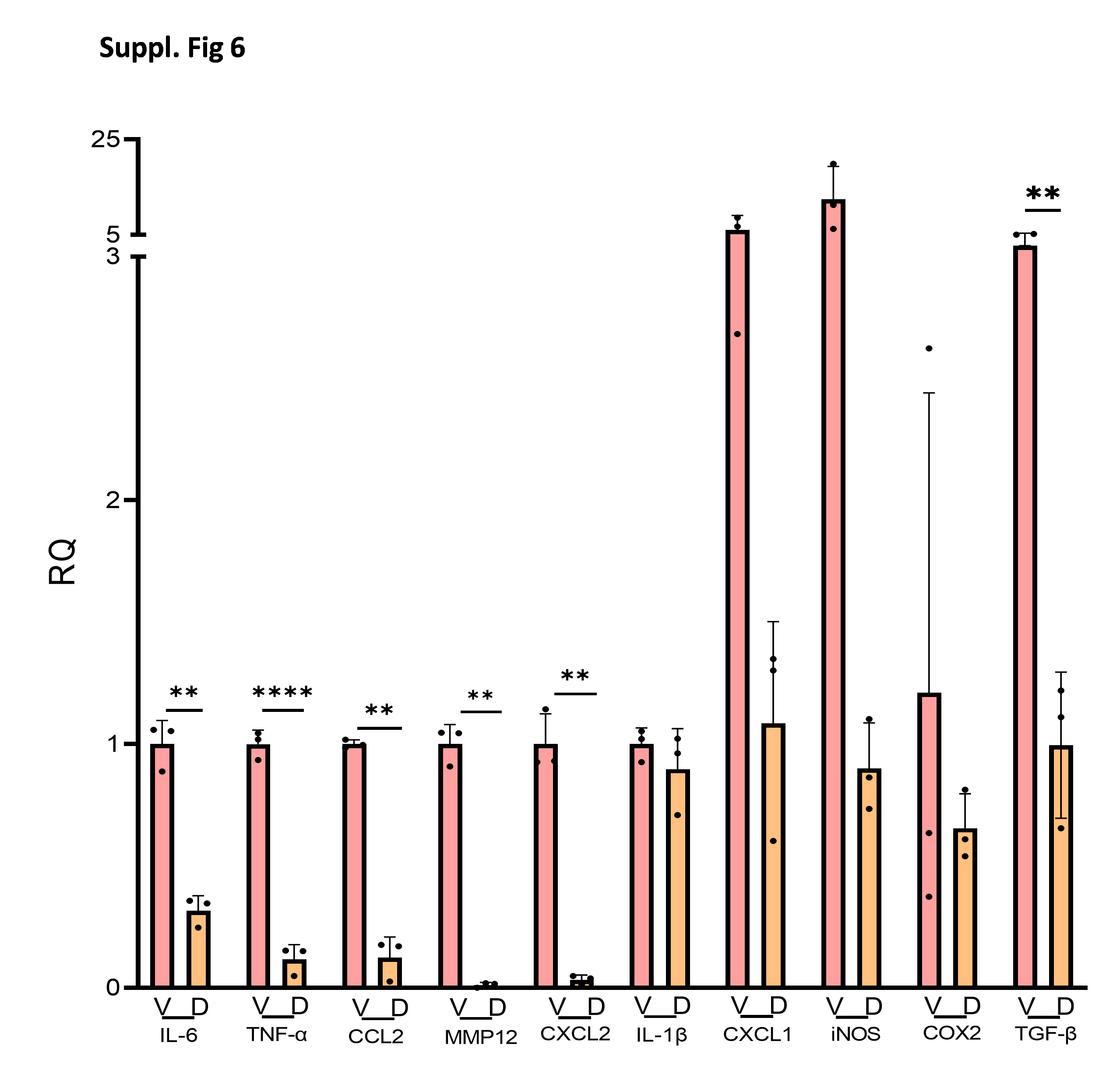

### Supplementary Fig. 7

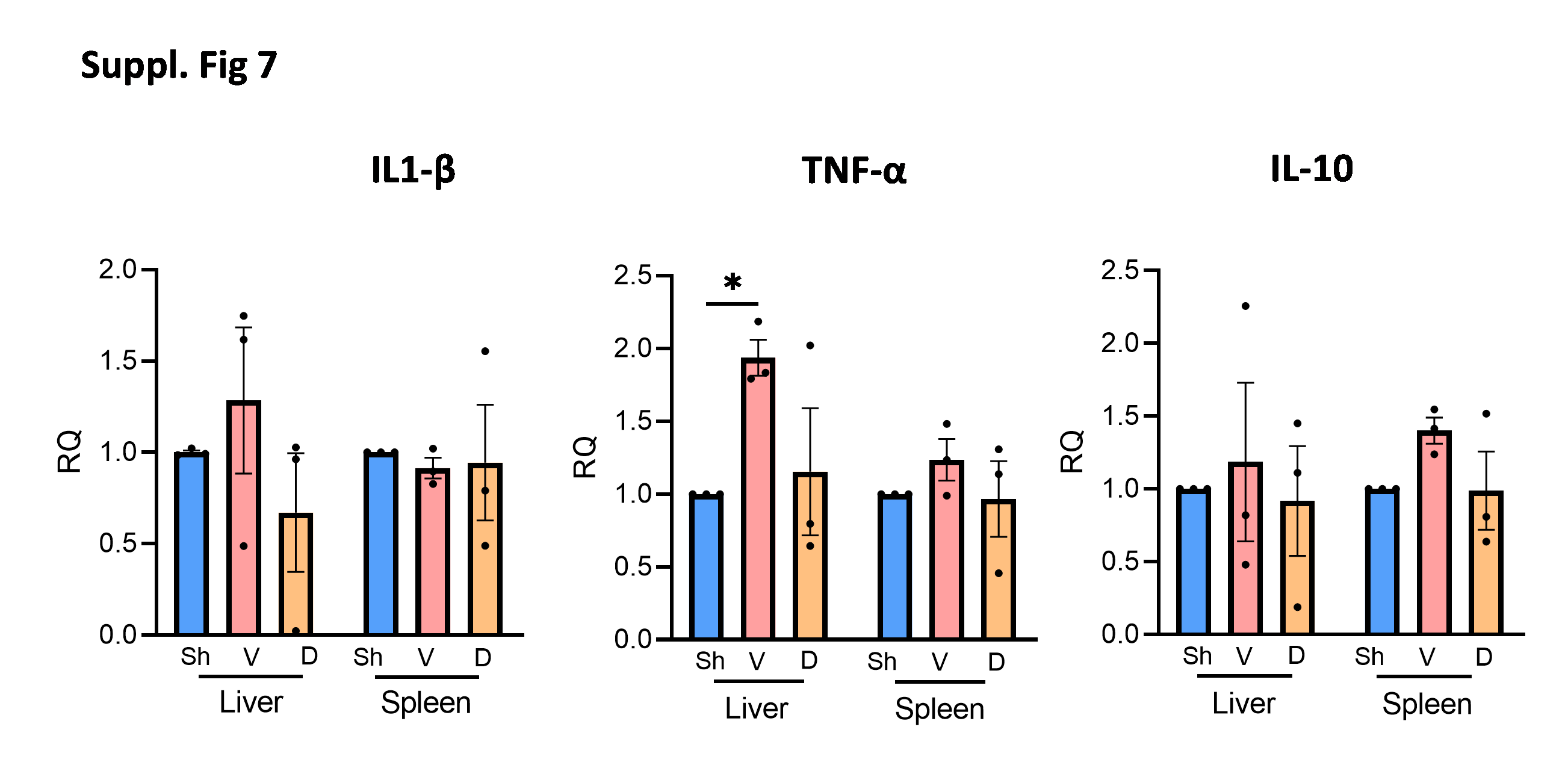

### Supplementary Fig. 8

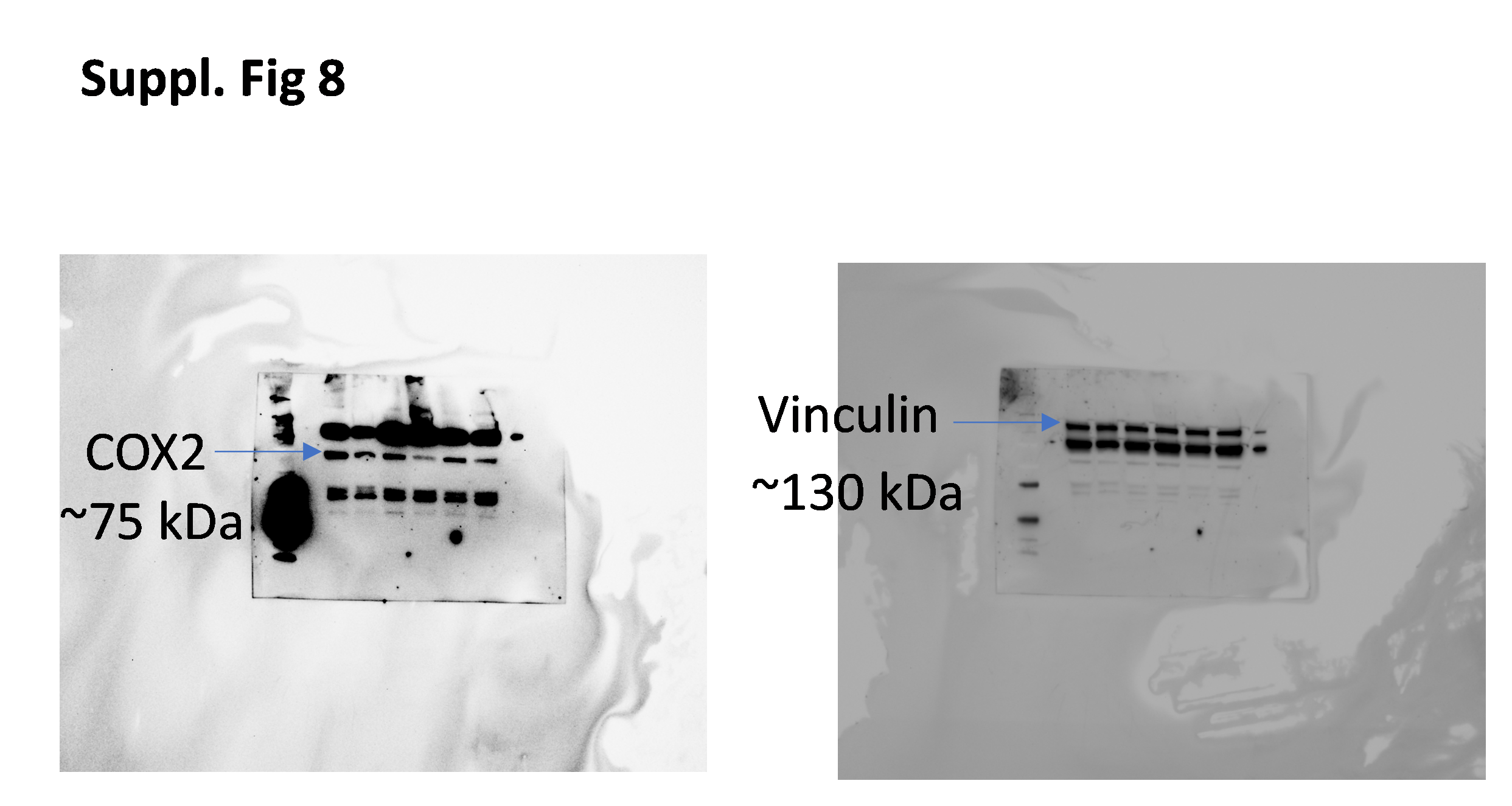
